# Involvement of denervated midbrain-derived factors in the formation of ectopic cortico-mesencephalic projection after hemispherectomy

**DOI:** 10.1101/2021.03.26.437124

**Authors:** Leechung Chang, Mayuko Masada, Masami Kojima, Nobuhiko Yamamoto

**Affiliations:** Laboratory of Cellular and Molecular Neurobiology, Graduate School of Frontier Biosciences, Osaka University, Suita, Osaka 565-0871, Japan; Biomedical Research Institute (BMRI), National Institute of Advanced Industrial Science and Technology (AIST), Ikeda, Osaka 563-8577, Japan; Core Research for Evolutional Science and Technology (CREST), Kawaguchi, Saitama 332-0012, Japan

## Abstract

Neuronal remodeling after brain injury is essential for functional recovery. After unilateral cortical lesion, axons from the intact cortex ectopically project to the denervated midbrain to compensate for the lost function, but the underlying molecular mechanisms remain largely unknown. To address this issue, we examined gene expression profiles in denervated and intact mouse midbrains after hemispherectomy at P6, when ectopic contralateral projection occurs robustly. The analysis showed that various axon growth-related genes were upregulated in the denervated midbrain, and most of these genes are reportedly expressed by astrocytes or microglia. To identify the underlying molecules, the receptors for candidate upregulated molecules were knocked out in layer 5 projection neurons in the intact cortex, using the CRISPR/Cas9-mediated method, and axonal projection from the knocked-out cortical neurons was examined after hemispherectomy. We found that the ectopic projection was significantly reduced when integrin subunit beta 3 (Itgb3) or neurotrophic receptor tyrosine kinase 2 (Ntrk2, also known as TrkB) was knocked out. Overall, the present study suggests that midbrain-derived glial factors whose expression is upregulated after hemispherectomy are involved in lesion-induced remodeling of the cortico-mesencephalic projection.

## Introduction

Neural circuits are reorganized after brain injury. One mechanism for the reorganization is compensation by new axonal growth into the denervated brain region from external sources. It is well known that compensatory axonal projections are formed after cortical lesion. Fundamentally, layer 5 neurons in the motor cortical area project ipsilaterally to the brain stem and contralaterally to the spinal cord. However, after a unilateral cortical lesion, descending axons from the intact cortex sprout and project contralaterally to the midbrain and hindbrain, and ipsilaterally to the spinal cord (Benowitz & Carmichael, 2010; Takahashi et al., 2009). Lesion-induced contralateral projection to midbrain nuclei such as the red nucleus is particularly well characterized (Tsukahara, 1981; Murakami & Higashi, 1988; Nah and Leong, 1976; Lee et al., 2004; Omoto et al., 2010). Functional compensation can also be achieved in the adult brain by this new projection but is insufficient for complete functional recovery. Therefore, enhancing the formation of axonal sprouting is one of the main strategies to overcome incomplete recovery after CNS injury.

Molecular manipulations have been proposed to facilitate the ectopic contralateral projection, by various methods such as nullifying the myelin-associated axon growth inhibitor NogoA or Nogo Receptor (Lee et al., 2004; Cafferty & Strittmatter, 2006), degrading CSPG (Starkey et al., 2012), or overexpressing growth-promoting transcription factors such as Stat3 (Lang et al., 2013) or Klf7 (Blackmore et al., 2012). On the other hand, the formation of compensatory circuits has been shown to be robust at infant developmental stages (Tsukahara, 1981; Kuang & Kalil, 1990; Omoto et al., 2010). Therefore, activation of intrinsic mechanisms may promote the ectopic projections and functional recovery. So far, trophic factors and axon growth regulatory components have been shown to contribute to compensatory connections in the spinal cord (Ueno et al., 2012; Fink et al., 2017). However, the molecular mechanisms are not fully understood. Moreover, the intrinsic mechanisms underlying formation of lesion-induced cortico-mesencephalic projections are completely unknown.

In the present study, we addressed this issue by hypothesizing that some attractive or promoting factors governing the formation of the lesion-induced ectopic contralateral projection are released from the denervated region. First, we investigated the time-course of the ectopic cortico-mesencephalic projections following hemispherectomy (the removal of one hemisphere) of juvenile mice. Next, lesion-induced gene expression in the midbrain was analyzed using RNA-seq. Finally, we attempted to identify the molecules that underlie the ectopic contralateral projection by means of specific gene knock-out using CRISPR/Cas9 and *in vivo* gene transfer methods.

## Results

### Time course of ectopic contralateral cortico-mesencephalic projections after hemispherectomy

First, we examined the time-course of ectopic contralateral cortico-mesencephalic projections after hemispherectomy. The ablation was performed at postnatal day 6 (P6), when ectopic projections are known to form robustly (Takahashi et al., 2009; Omoto et al., 2010). Axonal projections from the intact cortex were studied 2 to 7 days after hemispherectomy by implanting DiI crystals into the motor and somatosensory cortical areas (Figure 1A). Only two days after the hemispherectomy, many labeled axons were found on the opposite side of the midbrain, although a small number of cortical axons also crossed the midline even without ablation (Figure 1B). Sequential sections (200-μm thickness) showed that midline crossing occurs at certain levels of the midbrain, where the Edinger-Westphal nucleus (EW) and red nucleus (RN) exist (Supplementary Figure 1). This aspect was quantified in two sequential sections along the anterior–posterior axis by calculating the axon density index (see Materials and methods), which is the ratio of the axon density in the contralateral side midbrain to that in the ipsilateral side. The axon density indices were roughly 2-fold greater in both sections 2 days (P8) after hemispherectomy than those in the control without ablation (anterior: 0.063 ± 0.010 for control, 0.15 ± 0.025 for lesioned, p < 0.05; posterior: 0.059 ± 0.009, 0.11 ± 0.013, p < 0.05, Mann–Whitney *U* test) (Figure 1C). At later times, more labeled axons were found in the contralateral side. The axon density indices were approximately 4-fold (anterior: 0.054 ± 0.010, 0.23 ± 0.029, p < 0.05; posterior: 0.044 ± 0.009, 0.19 ± 0.030, p < 0.05, Mann–Whitney *U* test) and 5-fold (anterior: 0.050 ± 0.011, 0.25 ± 0.035, p < 0.05; posterior: 0.051 ± 0.005, 0.24 ± 0.030, p < 0.05, Mann–Whitney *U* test) greater at 4 days (P10) and 7 days (P13) after hemispherectomy, respectively (Figure 1B-C).

**Figure 1.**
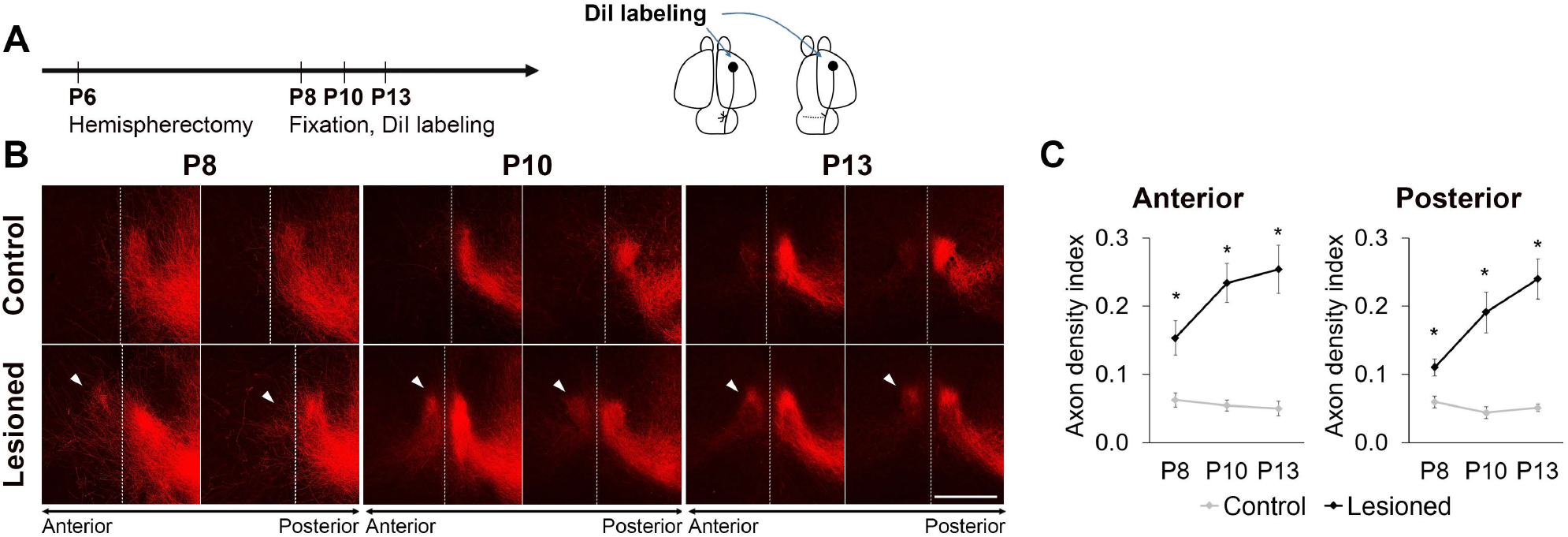
Midline-crossing cortico-mesencephalic axons increase after hemispherectomy. **A**, Schematic overview of the experiment. DiI was implanted into fixed brains to label cortical axons. **B**, Serial coronal sections showed that the contralateral projection to the midbrain starts to emerge only 2 days after surgery (P8) and increases at 4–7 days after surgery (P10, P13) compared with the control. Scale bar = 500 μm. Dashed lines indicate the midline. **C**, Quantitative analysis using the axon density index (see Materials and methods) of anterior and posterior sections (n = 4 for each group) confirmed a considerable increase in ectopic contralateral projection after hemispherectomy. * p < 0.05 against control; Mann–Whitney *U* test.

Because it is difficult, in the DiI-labeled fixed brain, to observe axons clearly at late developmental stages, perhaps due to diffusion of the dye in myelinated axons, the ectopic projections were further examined about one week after hemispherectomy by labeling cortical axons with a fluorescent protein. For this, cortical neurons in the motor or somatosensory area were transfected with an EGFP-expressing plasmid by *in utero* electroporation at E12.5, when layer 5 neurons are born (Figure 2A). In the control without ablation, EGFP-labeled axons invaded the midbrain and projected into and around the ipsilateral RN and EW, with few midline crossing axons, in accordance with the result of DiI labeling (Figure 2C, E). In contrast, a large number of labeled axons were found to project contralaterally after hemispherectomy. In the anterior midbrain, labeled axons accumulated in the vicinity of the EW on not only the ipsilateral but also the contralateral side (Figure 2B, B’). Furthermore, midline-crossing axons were present in a more dorsal region near the superior colliculus (SC) (Figure 2B’’). In the more posterior midbrain, labeled axons grew further into the ventrolateral part of the denervated midbrain (arrowheads in Figure 2D), and were distributed in the vicinity of the contralateral RN (Figure 2D). Importantly, after hemispherectomy, axonal accumulation in the contralateral midbrain was comparable to that on the ipsilateral side, and the contralateral projections were formed in a mirror image fashion of the ipsilateral projections.

**Figure 2.**
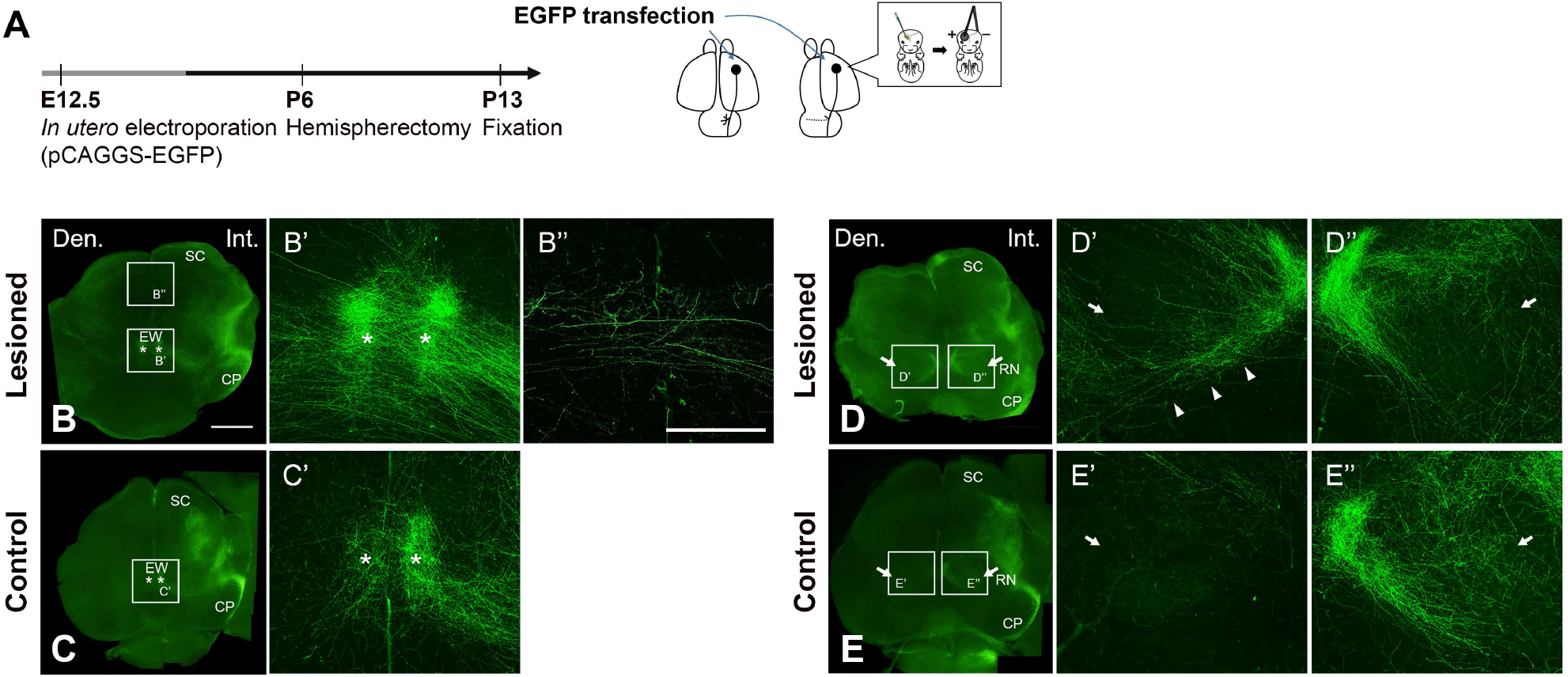
Contralateral cortical axons accumulate in midbrain nuclei 7 days after hemispherectomy. **A**, Schematic overview of the experiment. *In utero* electroporation-mediated transfection with *EGFP* was performed at E12.5 to label layer 5 cortical neurons. **B-E**, Coronal section images of hemispherectomized and control samples at P13 (7 days post-surgery). Left panels show low-magnification images of midbrain coronal sections, and right panels show magnified views of the indicated boxed areas of the left panels. After hemispherectomy, contralaterally projecting axons were found in the vicinity of the EW (asterisks), while only few midline-crossing axons were present in the control (**B, C**). Some of them were present more dorsally, near the SC. Bilateral projections to the RN (arrows) were formed after hemispherectomy in the posterior sections (**D, E**). Den., denervated side; Int., intact side. Scale bar = 1 mm (left panels), 500 μm (right panels).

To further elucidate the morphology of the lesion-induced projection, we utilized a tissue clearing method and sparse labeling of the cortical neurons using the Supernova system (Mizuno et al., 2014; Luo et al., 2016) to trace individual distinguishable axons (Supplementary Figure 2A). After hemispherectomy, contralaterally projecting axons were found to form branches just before the midline (Supplementary Figure 2B). Quantitative analysis showed that axon branching of contralaterally projecting axons occurred within 400 μm from the midline (Supplementary Figure 2C). These observations indicate that intact cortical axons form branches near the midline, project contralaterally at around 2 days after surgery, and continue to grow for several more days.

### Denervated midbrain promotes axon growth of deep layer cortical neurons

Next, our hypothesis that the denervated midbrain contains axon growth-promoting activity was tested *in vitro* by co-culturing a cortical slice (P1 motor cortex) with either denervated or normal midbrain tissue (Figure 3A). After one day in culture, axon elongation was examined by implanting DiI crystals into the fixed cortical slice. As shown in Figure 3B, axonal growth from the cortical explant was promoted when it was co-cultured with the denervated midbrain. The number of axons that grew to more than 300 or 400 μm from the ventricular edge was counted for quantification. The result revealed that the number was significantly larger when the explant was co-cultured with the denervated midbrain than with the normal midbrain (300 μm: 29.0 ± 7.5 for normal midbrain, 71.4 ± 10.6 for denervated midbrain, p = 0.0083; 400 μm: 14.6 ± 5.2, 47.9 ± 10.3, p = 0.019, Student’s *t*-test) (Figure 3C). Furthermore, retrograde labeling with DiI showed that projection neurons were mostly located in the area 400 μm to 600 μm from the pial surface, in which deep layer neurons are located (Figure 3D). Taken together, these results indicate that the denervated midbrain contains axon growth-promoting factors for deep layer projection neurons.

**Figure 3.**
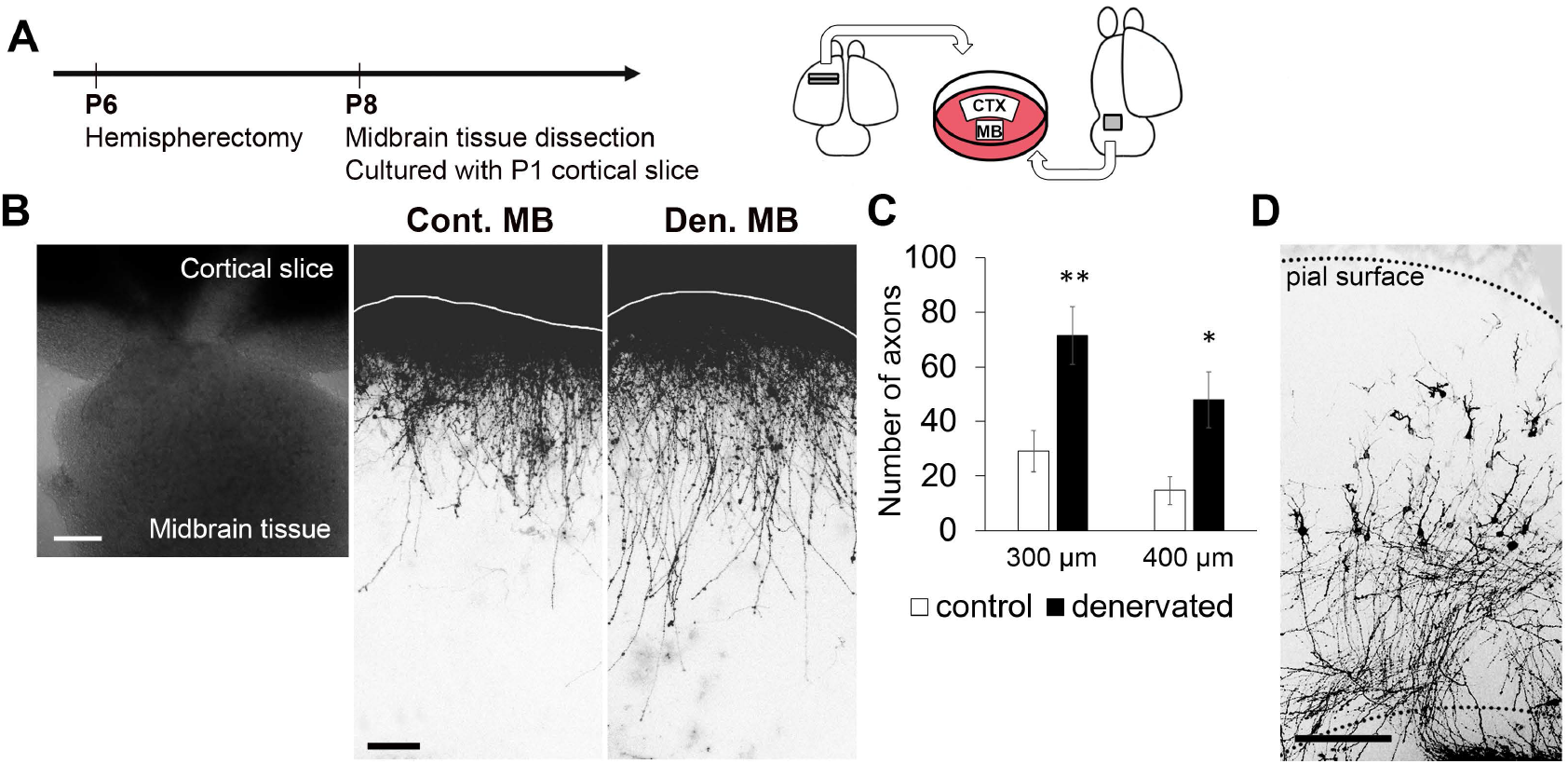
Co-culture with denervated midbrain promotes axon growth of deep layer cortical neurons. **A**, Schematic overview of the experiment. Co-culture of a cortical slice from intact P1 mice with a block of either the denervated midbrain two days after surgery or the same-aged midbrain without surgery. **B**, Representative images of co-culture (left), and labeled cortical axons (right) when cultured with control midbrain (Cont. MB) and denervated midbrain (Den. MB). The white line indicates the edge of the ventricular side of the cortical slice. Scale bar = 200 μm (left), 100 μm (right). **C**, Histogram showing the number of axons longer than 300 μm and 400 μm, measured from the ventricular edge, in co-cultures with control (n = 10) or denervated midbrain (n = 11). Co-cultures with denervated midbrain showed a greater number of longer cortical axons. * p < 0.05, ** p < 0.01 against control; Student’s *t*-test. **D**, Representative image of retrogradely labeled cells. Retrogradely labeled neurons were mostly found in the deep layers. Scale bar = 200 μm.

### RNA-seq analysis shows that glial cell-related genes are upregulated in the denervated midbrain

To investigate which genes are upregulated in the denervated midbrain, and thus candidate growth-promoting factors, we performed transcriptome sequencing and gene expression profiling. For this analysis, mRNAs were extracted from denervated midbrain 2 days (P8) and 4 days (P10) after hemispherectomy, when the ectopic projections are in the process of forming. These gene expression profiles were then compared with those of control midbrain and intact-side midbrain, and genes that were specifically upregulated or downregulated in the denervated midbrain were selected (FDR < 0.05, n = 2) (Figure 4A).

**Figure 4.**
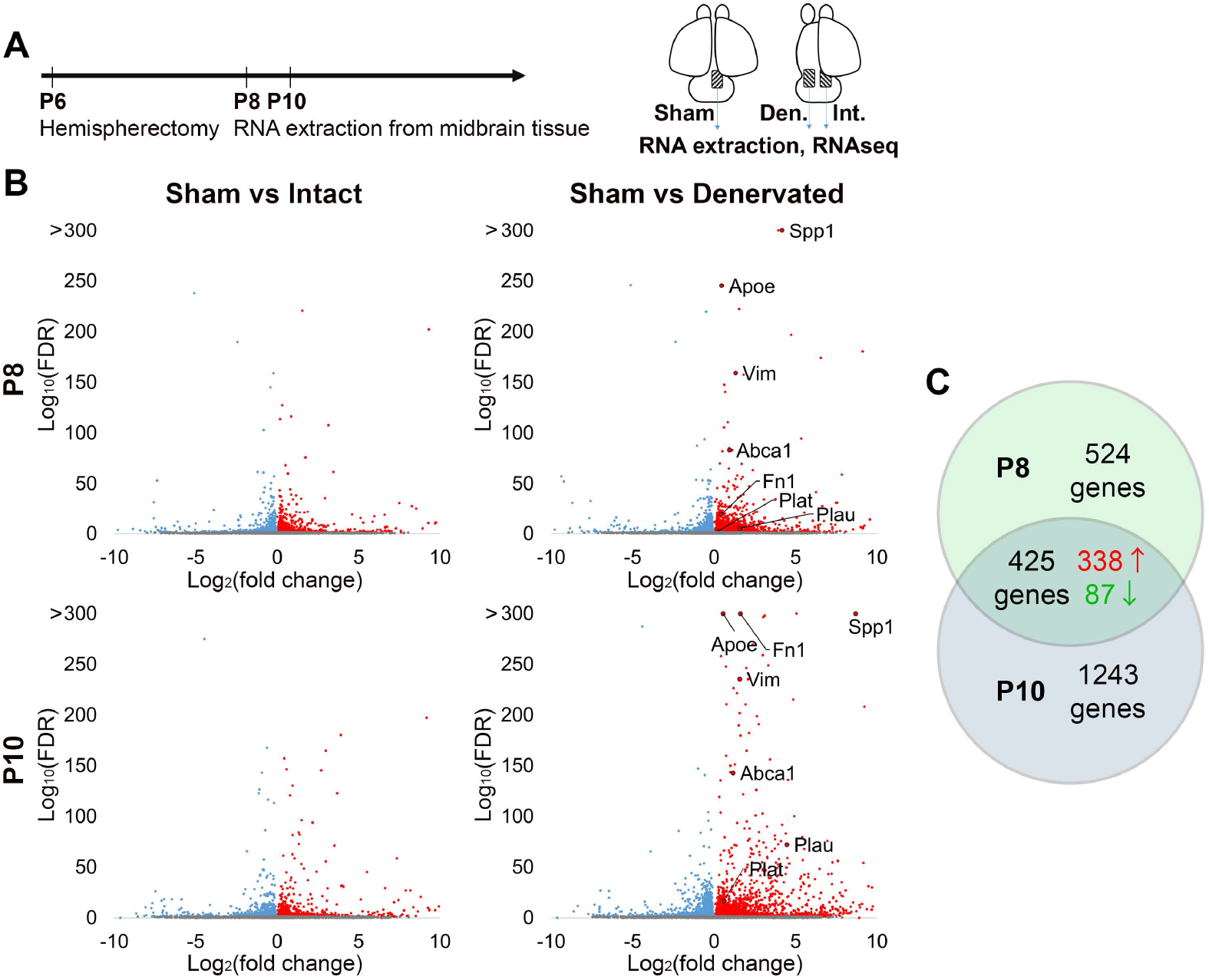
RNA-seq analysis shows that a subset of glial cell-derived genes is significantly upregulated in the denervated midbrain **A**, Schematic overview of the experiment. RNA-seq was performed using P8 or P10 (2 or 4 days post-surgery) midbrain taken from sham-operated and hemispherectomized mice (n = 2 for each group). **B**, Volcano plots comparing sham versus intact midbrain, and sham versus denervated midbrain. Differentially expressed genes (DEGs; FDR < 0.05) are highlighted in blue (downregulated) or red (upregulated). **C**, Venn diagram representation of DEGs in P8 (949 genes) and P10 (1668 genes). A total of 425 genes were common to the two stages (338 upregulated and 87 downregulated).

We found that 949 and 1668 genes were differentially regulated at P8 and P10, respectively, among which 338 upregulated genes and 87 downregulated genes were common to both stages (Figure 4B-C). We searched a public database of cell type-specific expression in mouse brain (https://www.brainrnaseq.org/; Zhang et al., 2014), and found that the majority of the upregulated genes were expressed by microglia and/or astrocytes (microglia: 250/338; astrocytes: 115/338). For example, complement protein genes (e.g., C1qa, C1qb) and phagocytosis-related genes (e.g., Fcgr3), which are known to be expressed by microglia, were upregulated, along with astrocyte-derived genes such as Vim and Serpina3n (Supplementary data 1). In support of this view, immunohistochemistry with antibodies against ionized calcium-binding adapter molecule 1 (Iba1) and glial fibrillary acidic protein (GFAP) demonstrated that many more reactive microglia and astrocytes were distributed in the denervated midbrain than on the opposite side (Supplementary Figure 3B-E). These results suggest that the upregulated genes are attributable to the glial cell response in the denervated midbrain.

The spatial distribution of upregulated genes was further investigated by performing *in situ* hybridization. Most analyzed genes were highly expressed in the vicinity of the cerebral peduncle (CP) of the denervated side, in which a large number of axons projecting to the midbrain and spinal cord degenerated after hemispherectomy (Figure 5). Some genes were rather restricted to the CP (e.g., Tyrobp, Spp1; Figure 5A–B), whereas others were broadly distributed along the ventral side of the denervated midbrain spreading to the midline (e.g., Fcgr3, Abca1, Vim, Fn1, Plat; Figure 5C–G). Thus, upregulated genes were confirmed to be expressed strongly on the denervated side.

**Figure 5.**
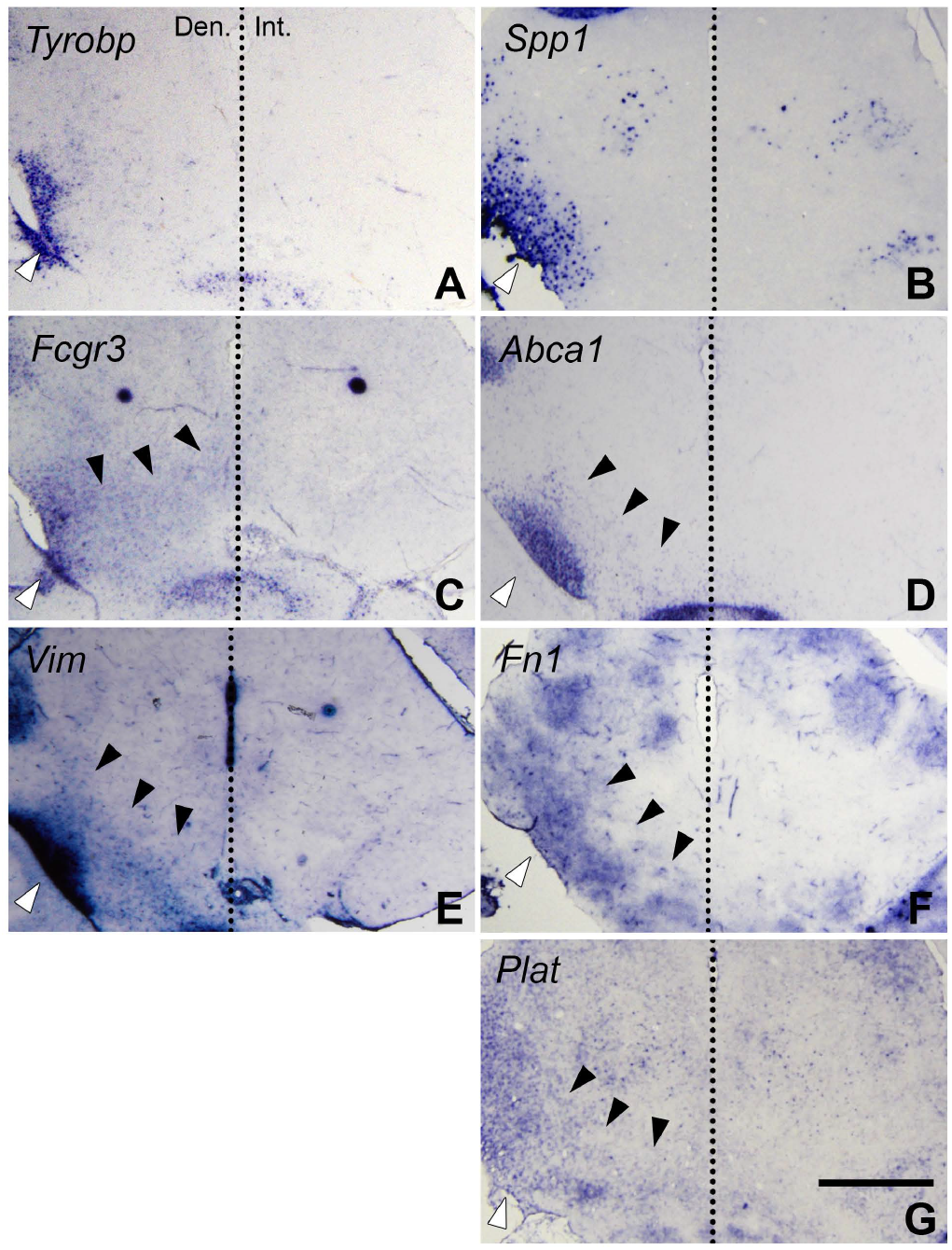
*In situ* hybridization confirms that lesion-induced genes are upregulated in the denervated midbrain. *In situ* hybridization was performed on P10 coronal sections of hemispherectomized mice. The expression of 7 representative upregulated genes is shown. The midline is indicated by dashed lines. White arrowheads indicate the CP of the denervated side. Black arrowheads indicate that some genes showed widespread signals in the region adjacent to the midline. Scale bar = 1 mm.

### Formation of ectopic contralateral projections is altered after CRISPR/Cas9-mediated KO of the receptors for the upregulated molecules

From the list of upregulated genes, those related with axon growth were selected as candidate regulators of lesion-induced axonal remodeling (Table 1). To identify the underlying molecules, we knocked out receptors for the above candidate molecules in layer 5 projection neurons. Furthermore, the receptors (e.g., TrkB, Igf1r) whose signaling pathway is potentially affected by the upregulated genes were also examined (see Table 1). For KO of these receptors, *in utero* electroporation-mediated transfection with vectors expressing Cas9 and sgRNAs was carried out together with the EGFP vector (Figure 6A). We found that the lesion-induced axonal projection in the denervated midbrain was markedly affected by KO of integrin subunit beta 3 (Itgb3) and neurotrophic receptor tyrosine kinase 2 (Ntrk2, also known as TrkB) (Supplementary Figure 4). Itgb3 is a receptor for extracellular molecules such as Spp1 and Fn1 (Humphries et al., 2006), both of which were upregulated in the denervated midbrain (Figure 5B, F, Supplementary Figure 6, Figure 6B). This receptor is expressed endogenously in layer 5 neurons (Supplementary Figure 5). TrkB is also expressed in cortical neurons (Supplementary Figure 5) and acts on BDNF signaling. In our RNA-seq data, the level of Bdnf mRNA was not elevated in the denervated midbrain (Figure 6B), but plasminogen activators (Plat, Plau), which can promote extracellular cleavage of proBDNF into the mature type (Pang et al., 2004), were highly increased in the denervated midbrain (Figure 5G, Figure 6B). Indeed, western blot analysis showed that mature BDNF was significantly more abundant in the denervated than the intact midbrain (ratio, 1.47 ± 0.11, p = 0.017, Student’s *t*-test) (Figure 6C), suggesting that proteolytic cleavage into mature BDNF is promoted in the denervated midbrain.

**Table 1.**
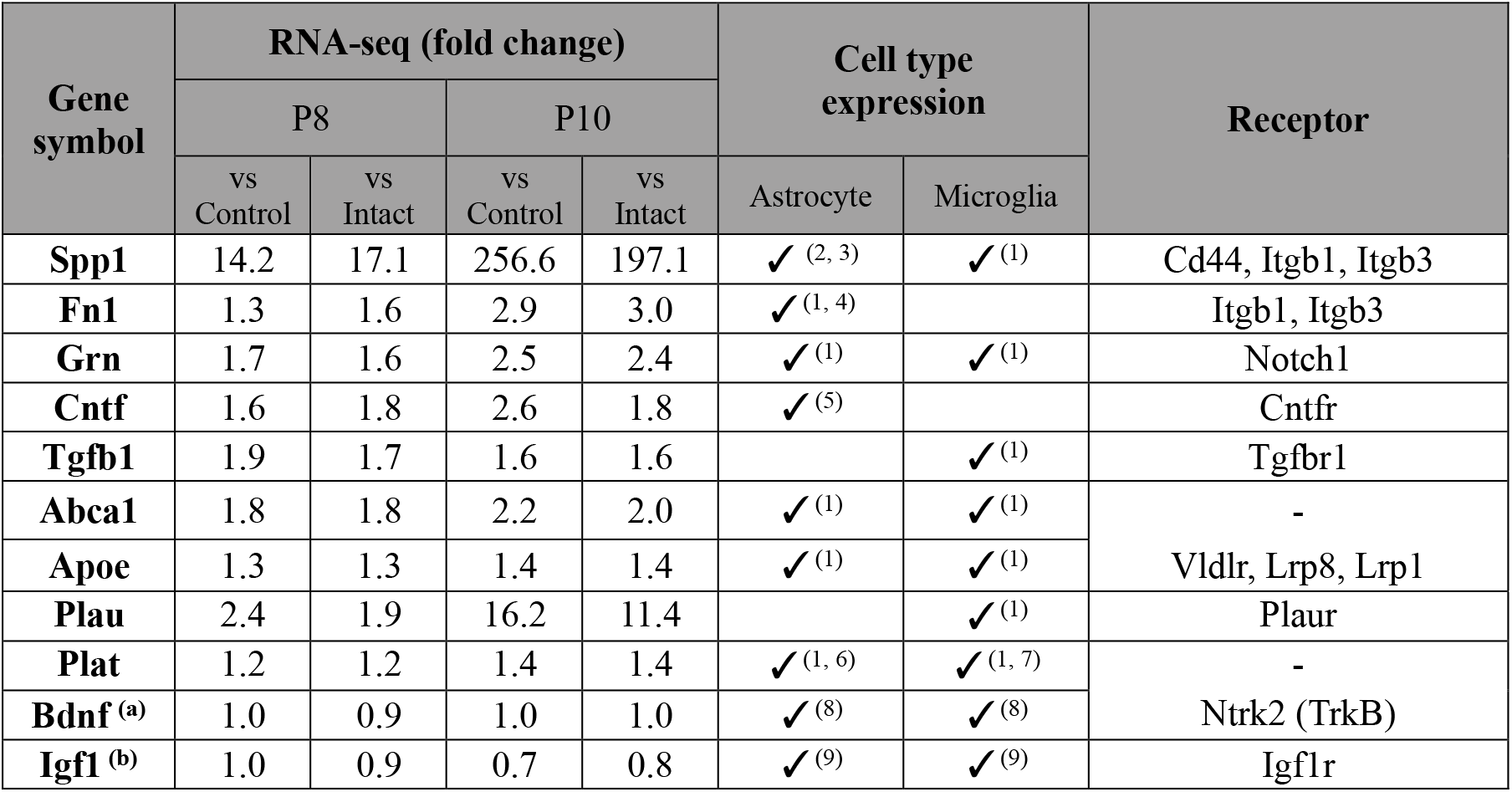
Upregulated genes related to axonal growth or regeneration. Their receptors are also presented in the table. Bdnf and Igf1 were not upregulated, but the signaling pathways of these genes may be affected by upregulated molecules. (a) Plau and Plat promote cleavage of proBDNF into mature BDNF (Pang et al., 2004). (b) Spp1 sensitizes corticospinal neuronal responses to IGF1, leading to an increase in the number of corticospinal sprouting neurons after cortical stroke (Liu et al., 2018). Cell type expression (astrocyte and microglia) of each gene is based on the following references: (1) Zhang et al., 2014; (2) Ellison et al., 1998; (3) Sinclair et al., 2005; (4) Tom et al., 2004; (5) Dallner et al., 2002; (6) Toshniwal et al., 1987; (7) Tsirka et al., 1995; (8) Dougherty et al., 2000; (9) Labandeira-Garcia et al., 2017.

**Figure 6.**
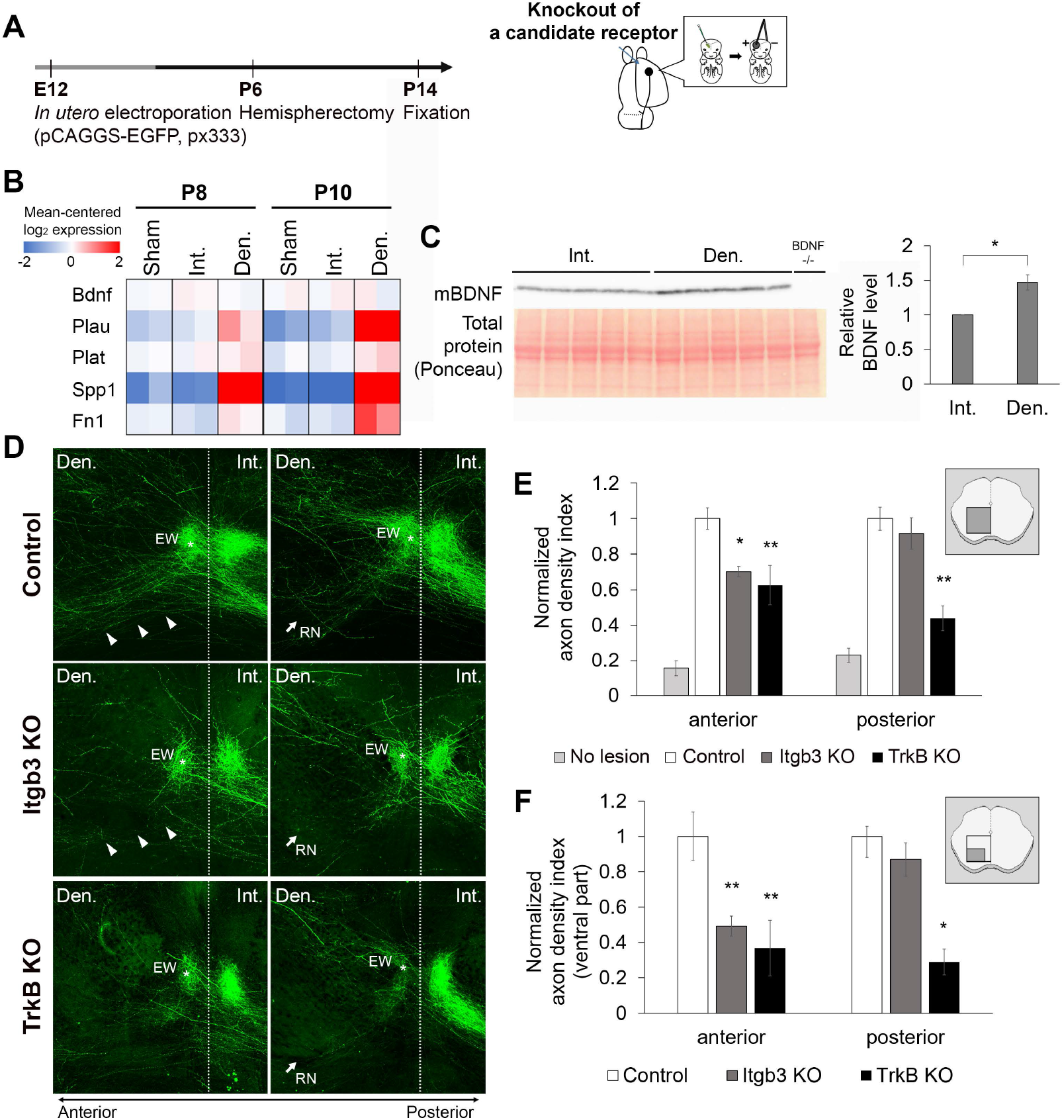
Formation of lesion-induced contralateral projection is impaired after KO of Itgb3 and TrkB. **A**, Schematic overview of the experiment. *In utero* electroporation was performed to knock out a specific receptor in layer 5 cortical neurons. **B**, Heat map of mean-centered log_2_ expression (FPKM) of the related genes. **C**, Western blot analysis shows a significant increase of mature BDNF amount in the denervated-side midbrain (n = 5) compared with the intact side (n = 5) taken from P10 mice (4 days post-surgery). * p < 0.05; Student’s *t*-test. **D**, After hemispherectomy, ectopic axons projecting from Itgb3 KO and TrkB KO neurons were reduced, compared with the control (without KO; after hemispherectomy). Anterior and posterior sections were used for the analysis. Dashed lines indicate the midline. Note that axon projection to the ventrolateral portion is impaired in Itgb3 KO (arrowheads). Scale bar = 500 μm. **E**, Normalized axon density indices (axon density index normalized by that for without KO) were calculated in the anterior and posteriors sections for each case (no lesion, n = 6; control, n = 7; Itgb3 KO, n = 10; TrkB KO, n = 6). **F**, Normalized axon density indices were similarly calculated in the ventrolateral region. The region of interest for each analysis (E, F) is indicated by the illustrations on the right. * p < 0.05, ** p < 0.01 against control; one-way ANOVA with Tukey’s *post hoc* test.

Quantitative analysis of the two sequential sections demonstrated that lesion-induced axonal projection was reduced by KO of these two receptors (Figure 6D-F, Supplementary Figure 7). In Itgb3 KO samples, ectopic contralateral projections were reduced in the anterior section (Figure 6D). Indeed, the axon density index in the anterior section was significantly lower in Itgb3 KO (axon density index normalized by that for without KO, 0.70 ± 0.028, p = 0.040, one-way ANOVA with Tukey’s *post hoc* test), but not in the posterior section (normalized axon density index, 0.92 ± 0.088, p = 0.90) (Figure 6E). In particular, axons projecting further into the ventrolateral part of the denervated midbrain, where the expression of Spp1 and Fn1 was increased after hemispherectomy (Figure 5B, F, Supplementary Figure 6), were greatly reduced in the anterior section (Figure 6D). Quantitative analysis of the restricted area including the ventrolateral part showed a marked decrease in the axon density index (normalized axon density index, 0.49 ± 0.057, p = 0.0082, one-way ANOVA with Tukey’s *post hoc* test) (Figure 6F). The knocked-out phenotype was also examined for the beta-1 subtype (Itgb1), which is also expressed in layer 5 neurons (Nieuwenhuis et al., 2018), but no reduction in contralateral axonal projection was found after hemispherectomy (Supplementary Figure 4). On the other hand, TrkB KO samples showed a substantial decrease in the axon density index after hemispherectomy in both anterior and posterior sections, with a more profound reduction in the posterior section (normalized axon density index, 0.63 ± 0.11 for anterior, p = 0.0072; 0.44 ± 0.070 for posterior, p = 0.0018, one-way ANOVA with Tukey’s *post hoc* test) (Figure 6D, E). These results indicate that Itgb3 and TrkB KO reduced ectopic axonal projections in the denervated midbrain in a distinct fashion after hemispherectomy.

## Discussion

The present morphological and molecular expression analyses demonstrate that glial cell-related axon growth-promoting factors are expressed in the denervated midbrain when robust remodeling of the cortico-mesencephalic projection takes place after hemispherectomy. Furthermore, CRISPR/Cas9-mediated functional analysis demonstrate that these factors are involved in the remodeling of the the cortico-mesencephalic projection. In particular, KO of Itgb3 or TrkB suppressed the formation of ectopic contralateral projections. Thus, the present study revealed for the first time that these specific molecules contribute to remodeling of the cortico-mesencephalic projection after hemispherectomy.

### The cortico-mesencephalic projection is rapidly remodeled after hemispherectomy of juvenile mice

A morphological experiment with axon tracing showed that remodeling of the cortico-mesencephalic projection took place after hemispherectomy, in accordance with previous reports (Tsukahara, 1981; Murakami & Higashi, 1988; Nah and Leong, 1976; Lee et al., 2004; Takahashi et al., 2009; Omoto et al., 2010). Robust formation of the ectopic contralateral projection is consistent with previous findings that axonal sprouting is extensive in young animals (Tsukahara, 1981; Kuang & Kalil, 1990; Omoto et al., 2010). Moreover, our results demonstrated that only a few days after hemispherectomy, axons from the intact cortex invaded the denervated midbrain. This rapid formation of the ectopic contralateral projection raised the possibility that axonal sprouting occurs in the regions near the midline, which was supported by the observation that contralaterally projecting axons originated from branch points adjacent to the midline (Supplementary Figure 2). Furthermore, the subsequent increase in the ectopic contralateral projection is likely due to an increase in the number of midline-crossing axons (Lee et al., 2004) rather than extensive branch formation in the small number of pre-existing contralateral axons that exist even in intact animals.

### Axon growth-promoting factors expressed in the denervated midbrain are involved in lesion-induced remodeling

Two previous studies have shown transcriptome profiles of the denervated region (Bareyre et al., 2002; Kaiser et al., 2019), but they could not lead to the identification of specific molecules that are actually involved in axonal sprouting and remodeling. Our CRISPR/Cas9-mediated KO study demonstrated that two distinct molecular mechanisms contribute to the formation of the ectopic projections. First, Itgb3 KO showed a significant decrease of the ectopic projections in the ventrolateral portions of the midbrain where Spp1 and Fn1 are upregulated (Figure 5B, F, Supplementary Figure 6) (Tom et al., 2004; Myers et al., 2011; Wright et al., 2014; Duan et al., 2015). It is plausible that developmental mechanisms guide axons to their proper targets in both intact and denervated midbrain, as the lesion-induced contralateral projection resembles the intact ipsilateral projection (Figure 2D). The axonal growth effect of Spp1 and Fn1 may enhance this process, helping axons to reach their targets. Second, TrkB KO showed a striking decrease of the ectopic projections, which may be due to non-utilization of mature BDNF, whose concentration should be elevated by proteolytic cleavage in the denervated side (see below, Figure 6C). BDNF–TrkB signaling may also contribute to the final connections with target cells, as it is known to promote axonal branch formation (Cohen-Corey et al., 2010; Granseth et al., 2013). The fact that the ipsilateral projection was hardly affected by knocking out Itgb3 and TrkB indicates that these mechanisms may not contribute to overall axonal growth of the cortico-mesencephalic projection.

Previous work has demonstrated that blocking BDNF–TrkB signaling hinders the lesion-induced sprouting of the corticospinal projection (Ueno et al., 2012), in which Bdnf expression was not upregulated in the denervated spinal cord after the cortical lesion. The present results also show that the Bdnf mRNA level was unchanged in the denervated midbrain, but that the level of mature BDNF protein was upregulated (Figure 6C). It is known that proBDNF is released extracellularly (Yang et al., 2009), and post-transcriptional modification of proBDNF mediated by plasminogen activators has been implicated in various neuronal functions, such as long-term potentiation and long-term memory (Pang et al., 2004; Kojima et al., 2020b). Thus, we speculate that plasminogen activators upregulated in the denervated midbrain promote the extracellular cleavage of proBDNF and production of mature BDNF.

### Glial cells as the source of the axon growth-promoting factors

Which cell types produce these axon growth-promoting factors? Our RNA-seq analysis showed that many glial cell-related genes were upregulated in the denervated midbrain after hemispherectomy, in accordance with previous investigations of corticospinal projection (Tsujioka and Yamashita, 2019; Kaiser et al., 2019). Furthermore, the upregulated genes Spp1 (Ellison et al., 1998; Sinclair et al., 2005), Fn1 (Tom et al., 2004), and plasminogen activators (Toshniwal et al., 1987; Tsirka et al., 1995) have been shown to be expressed by glial cells (Supplementary Figure 6).

We have also observed that reactive glial cells emerged in the denervated midbrain (Supplementary Figure 3B–E), and their distribution was similar to that of the upregulated genes. Furthermore, lesion-induced ectopic axons ran in the region where reactive astrocytes reside (Supplementary Figure 3B’, C’). This further supports the view that reactive glial cells express the molecules responsible for the circuit remodeling.

After CNS injury, a glial scar is formed at the lesion site, which is known to be inhibitory to axon regeneration (Silver and Miller, 2004). It is likely that reactive glial cells in the denervated midbrain have different properties from the scar-forming glial cells, as the denervated midbrain is distant from the lesion site. As in the case of “alternatively” activated glial cells (Hu et al., 2015; Liddelow and Barres, 2017), glial cells in the denervated midbrain may act positively on axonal growth.

### Other possible mechanisms for lesion-induced remodeling

In the present study, impairing Itgb3- and TrkB-mediated signaling did not block sprouting completely, which implies that other mechanisms also operate in this process. The expression of axon growth-inhibitory molecules may decrease after hemispherectomy, as manipulations that remove inhibitory molecules such as Nogo or CSPG can facilitate axonal sprouting (Cafferty and Strittmatter, 2006; Starkey et al., 2012). However, we could not find any decline of these inhibitory molecules from our RNA-seq result. Another possibility is that a midline barrier exists which prevents axon growth, similar to the Ephrin-B3/EphA4 signaling pathway in the spinal cord (Kullander et al., 2001; Yokoyama et al., 2001). However, this is also unlikely, as the cortico-mesencephalic projection has been shown to be normal in these KO mice (Yokoyama et al., 2001; Serradj et al., 2014).

The involvement of neuronal activity has to be considered, because it has been reported that neuronal activity promotes axonal branching (Uesaka et al., 2005). Indeed, lesion-induced sprouting in the corticospinal tract is enhanced by electrical stimulation of the motor cortex (Carmel et al., 2010; Carmel and Martin, 2014) and rehabilitation (van den Brand et al., 2012; Wahl et al., 2014), which led to functional recovery. The expression of sprouting-inducing factors and/or receptor molecules may also be regulated in an activity-dependent fashion (Yap and Greenberg, 2018).

To date, a model in which axonal regeneration and/or sprouting after brain lesion is enhanced by removal of growth-inhibitory factors has been emphasized (Lee et al., 2004; Cafferty & Strittmatter, 2006; Starkey et al., 2012). On the other hand, the present study demonstrates a contribution of endogenous growth-promoting factors, whose expression is robust at early developmental stages. Therefore, molecular manipulations that enhance such an attractive mechanism may also be effective for axonal remodeling, and may provide a novel insight into new therapeutic strategies for functional recovery.

## Materials and methods

### Animals

ICR mice (Japan SLC) were used in this study. All experiments were conducted under the guidelines for laboratory animals of the Graduate School of Frontier Biosciences, Osaka University. The protocol was approved by the School’s Animal Care and Use Committee.

### Hemispherectomy

At postnatal day 6 (P6), mice were anesthetized with isoflurane (Wako). After cutting the skin, a small window (about 2 mm x 2 mm) was made using micro-scissors on the right side of the skull. The right hemisphere was completely removed using an aspirator. After the ablation, the pups recovered on a heating pad and were then returned to their mother. For sham operation, only the skin was cut.

### Anterograde axonal labeling using 1,1′-dioctadecyl-3,3,3′,3′-tetramethylindocarbocyanine perchlorate (DiI)

Mice were sacrificed two, four or seven days after hemispherectomy, and whole brains were fixed in 4% paraformaldehyde (PFA) in 0.1 M phosphate buffer (PB, pH 7.4). DiI crystals (Invitrogen) were implanted into the motor cortical areas of hemispherectomized and control brains. After incubation for 4 to 5 weeks in 4% PFA at 37°C, 200-μm thickness coronal sections were cut by a vibratome (DTK-1000; Dosaka). The sections were mounted in phosphate-buffered saline (PBS) under coverslips for microscopic observation (see below).

### Axon labeling by *in utero* electroporation

*In utero* electroporation was performed on E12.5 mouse embryos, according to previous studies (Fukuchi-Shimogori and Grove, 2001; Saito and Nakatsuji, 2001; Tabata and Nakajima, 2001). In brief, a pregnant mouse was anesthetized with isoflurane (Wako) using inhalation anesthesia equipment (KN-1071-1; Natsume). Approximately 1 μl of plasmid solution was injected into the lateral ventricle of the left hemisphere using a glass capillary and an injector (IM-30; Narishige). Cortical axons were labeled with pCAGGS-EGFP (0.5 μg/μl) (Hatanaka and Murakami, 2002), and electrical pulses (37 V with five 50-ms pulses at intervals of 950 ms) were then applied with tweezer electrodes (LF650P1; BEX) connected to an electroporator (CUY21; BEX).

Hemispherectomy was performed at P6 in these electroporated mice. After surgery, they were deeply anesthetized and perfused with PBS followed by 4% PFA in 0.1 M PB. To visualize axons, the brains were post-fixed with the same fixative overnight, and 200-μm thickness coronal sections were cut by a vibratome (DTK-1000; Dosaka). The sections were permeabilized with 1% Triton X-100 for 1 h at room temperature (RT), and blocked with blocking solution (5% normal goat or donkey serum, 0.3% Triton X-100 in PBS) for 1 h at RT. They were incubated with rat monoclonal anti-GFP (1:2000, GF090R; Nacalai Tesque) in the blocking solution at 4°C overnight. After extensive washes, the sections were incubated with appropriate secondary antibodies at 4°C overnight. After washing, the slices were mounted in medium containing 2.3% DABCO, 1 μg/ml DAPI and 50% glycerol for microscopic observation.

### CRISPR/Cas9–mediated gene KO in cortical neurons

To knock-out specific genes, a plasmid containing humanized Cas9 and dual single-guide RNAs was made as previously described (px333; Maddalo et al., 2014), based on pX330-U6-Chimeric_BB-CBh-hSpCas9 (Cong et al., 2013). pX330-U6-Chimeric_BB-CBh-hSpCas9 was a gift from Feng Zhang (Addgene plasmid # 42230; http://n2t.net/addgene:42230; RRID: Addgene_42230). The sequence of single-guide RNAs (sgRNAs) for each gene (Supplementary Table 2) was chosen from the Brie sgRNA library (Doench et al., 2016). For genes that were not included in the database, sgRNAs were designed using CHOPCHOP (http://chopchop.cbu.uib.no/; Labun et al., 2019). The px333 plasmid designed to target a specific gene (3.5 μg/μl) was electroporated together with pCAGGS-EGFP (0.5 μg/μl) into the E12.5 mouse brain by *in utero* electroporation (see above). Transfection of with either the empty px333 vector or with pCAGGS-EGFP alone was performed as a control. After electroporation, hemispherectomy and visualization of cortico-mesencephalic axons were performed as described above.

### Validation of KO efficiency

To validate KO efficiency, a primary cortical neuron culture was prepared as described previously (Kitagawa et al. 2017). After *in utero* electroporation at E12.5 (see above), pregnant mice were anesthetized again at E15 with pentobarbital (50 mg/kg, i.p.), and the GFP-positive area of the cerebral cortex was dissected from embryos in ice-cold Hanks’ balanced salt solution. The tissue was then minced with fine scissors and incubated with 0.125% Trypsin-EDTA (Gibco) for 5 min and dissociated thoroughly by pipetting. After a brief centrifugation, the cells were resuspended in DMEM/F12 medium (Life Technologies) supplemented with 10% fetal bovine serum and plated in 4-well culture dishes (Thermo Scientific) coated with 0.1 mg/ml poly-L-ornithine (Sigma-Aldrich). The cultures were maintained at 37°C in an environment of 5% CO_2_ and humidified 95% air, and were fixed after 2 days in 4% PFA in PBS at RT for 10 min. The cells were immunostained with rat anti-GFP (1:2000, GF090R; Nacalai Tesque) and goat anti-TrkB (1:250, AF1494; R&D Systems). Cells were extensively washed and incubated with appropriate secondary antibodies at 4°C overnight. Images were captured with a fluorescence microscope (IX71 with 10× or 20× objective lens; Olympus), and analyzed with ImageJ to measure fluorescence intensity.

Alternatively, Neuro2a cells were used for the validation. The cells were cultured in high glucose DMEM (Thermo Fisher Scientific) with 10% FBS, and transfected with plasmids (pCAGGS-EGFP and the px333 plasmids) using Lipofectamine 2000 (Thermo Fisher Scientific) according to the manufacturer’s instructions. The culture was maintained until DIV 7, and the cell lysates were subjected to western blot analysis (see below) to evaluate the expression level of the target gene. To estimate KO efficiency, the transfection efficiency (the ratio of the number of EGFP-positive cells to the total number of cells) was taken into account.

### Sparse labeling of axons and tissue clearing

To label axons sparsely, the Supernova system was used (Luo et al, 2016; Mizuno et al, 2014). Supernova vectors pTRE-Flpe-WPRE (pK036) and pCAG-FRT-stop-FRT-tRFP-ires-tTA-WPRE (pK037) were kindly gifted from Dr. T. Iwasato, National Institute of Genetics, Japan. These vectors were co-transfected with pCAGGS-EGFP by *in utero* electroporation as described above. After surgery (see above), mice were perfused with PBS and 4% PFA in 0.1 M PB. The brain was post-fixed overnight, and 1-mm coronal sections of the midbrain were cut by a vibratome (DTK-1000; Dosaka). Tissue clearing of these thick sections was performed using the SeeDB2 method as previously described (Ke et al., 2016). Briefly, sections were serially incubated in solutions with increasing concentrations of Omnipaque 350 (Daiichi-Sankyo), which contains 2% Saponin (Nacalai Tesque). Sections were imaged by confocal microscopy (TCS-SP5; Leica), and tiled images of the entire sections were acquired with a 10× objective lens in 10-μm steps. Axons were traced using Simple Neurite Tracer, a plug-in for ImageJ.

### Anterograde and retrograde labeling in organotypic cortical slice cultures

Organotypic cortical slice cultures were prepared as described previously (Yamamoto et al., 1989; Yamamoto et al., 1992). In brief, about 300-μm-thick coronal slices were dissected from P1 mouse cortex. A block of the midbrain (approximately 1 mm × 1 mm x 0.3 mm) was dissected near the midline and ventral to the cerebral aqueduct from P8 brain with and without hemispherectomy. A cortical slice and the midbrain block were placed on collagen gel with DMEM/F-12 (Invitrogen) supplemented with 10% fetal bovine serum (Hyclone). To make collagen gel, rat tail collagen was mixed with 10× DMEM/F-12 (Gibco) at a ratio of 9:1 with one volume of 0.25% sodium bicarbonate. The mixed solution was put on a cell culture insert (Millicell-CM, PICMORG50; Millipore) and allowed to harden. The cultures were maintained for 24–48 h at 37°C in an environment of 5% CO_2_ and humidified 95% air.

DiI crystals were implanted into the cortical slice for anterograde labeling or into the midbrain explant for retrograde labeling. DiI-labeled axons were imaged by confocal microscopy (ECLIPSE FN with EZ-C1; Nikon), and stack images were acquired with a 10× objective lens in 5-μm steps. Using ImageJ software, the number of axons crossing a line set to either 300 or 400 μm from the ventricle edge was counted.

### RNA-seq analysis

Midbrain tissue was collected from hemispherectomized and sham-operated mice at P8 and P10. Total RNA was extracted from the tissue using an RNeasy Plus Mini Kit (QIAGEN), and sequencing was performed on an Illumina HiSeq 2500 (BGI Japan). Sequenced reads were mapped to the mouse reference using Bowtie2 (Langmead et al., 2012), and gene expression level was calculated with RSEM software (Li et al., 2011). Differentially regulated genes were defined by false discovery rate (FDR) < 0.05 (n = 2). To explore cell type-specific expression profiles of upregulated genes, a public database (www.brainrnaseq.org; Zhang et al., 2014) was used, and genes with fragments per kilobase of transcript per million mapped reads (FPKM) > 10 were defined as being expressed by that cell type.

### *In situ* hybridization

*In situ* hybridization was performed as previously described (Liang et al., 2000; Zhong et al., 2004). To prepare RNA probes, cDNA was synthesized from P8 mouse total RNA, and the DNA fragments of genes of interest were amplified by PCR with a pair of primers (Supplementary Table 1). The primer pairs were mostly designed based on the Allen Mouse Brain Atlas (Lein et al., 2007). Each cDNA fragment was then cloned into the pGEM-T vector. To produce linearized templates, the inserts were PCR-amplified with primers containing T7 and SP6 promoter sequences (TTGTAAAACGACGGCCAGTG and TGACCATGATTACGCCAAGC). DIG-labeled RNA probes were then synthesized (DIG RNA Labeling Mix, Roche) following the manufacturer’s instructions.

Mouse brains were fixed with 4% PFA in 0.1 M PB at 4°C and cryopreserved with 30% sucrose in PBS. The brains were sectioned into 20-μm-thick coronal sections using a cryostat. The sections were subjected to re-fixation and acetylation. After pre-hybridization, they were hybridized with the DIG-labeled probe (4 μg/ml) at 60°C. After washing, the sections were incubated with alkaline phosphatase (AP)-conjugated anti-DIG antibody (1:1000, 11093274910; Roche) at 4°C for two days. Finally, the hybridized probes were visualized with AP substrate (BM Purple, Roche) at RT.

### Immunohistochemistry

For immunostaining of various molecules, the brains were perfused in 4% PFA in 0.1 M PB and post-fixed with the same fixative for 4 h or overnight. They were then equilibrated with 30% sucrose in PBS, frozen in OCT compound (Sakura Finetech), and sectioned at 20 or 50 μm using a cryostat (CM1850; Leica Microsystems). The sections were permeabilized with 0.1% Triton X-100 for 1 h at RT and blocked for 1 h at RT in blocking solution. They were incubated with the following primary antibodies in blocking solution at 4°C overnight: mouse monoclonal anti-GFAP (1:1000, G3893; Sigma-Aldrich), rabbit polyclonal anti-GFAP (1:100, G9269; Sigma-Aldrich), rabbit polyclonal anti-Iba1 (1:500, 019-19741; Wako), goat polyclonal anti-Spp1 (1:50, AF808; R&D Systems), and rabbit anti-Plat (1:100, ASMTPA-GF-HT; Molecular Innovations). Sections were washed, incubated with appropriate secondary antibodies at 4°C overnight, and mounted.

### Western blot analysis

Midbrain tissues from the denervated and intact sides were rapidly collected from hemispherectomized mice at P10 (4 days after the operation). The dissected tissues were homogenized in RIPA buffer containing a protease inhibitor cocktail (P8340; Sigma-Aldrich), and the supernatants were collected after centrifugation at 16,000 × *g* for 30 m. The protein concentrations of the supernatants were determined using a BCA Protein Assay Kit (Thermo Fisher Scientific). The same amount of protein (25 μg) of each sample was applied to SDS-PAGE using 15% polyacrylamide gels in the presence of β-mercaptoethanol. After electrophoresis, proteins were transferred to PDVF membrane (Bio-Rad) using a Mini Trans-Blot Cell (Bio-Rad). The membrane was blocked with Tris-buffered saline containing 0.1% Tween-20 (TBS-T) and 5% skim milk (Nacalai Tesque) for 1 h at RT, and then incubated with mouse anti-BDNF antibody (1:1000, 327-100; Icosagen) in 2% skim milk in TBS-T at 4°C overnight (Kojima et al., 2020a). Primary antibody was detected by incubating the membrane with peroxidase-conjugated anti-mouse IgG antibody (1:10000, 115-035-146; Jackson ImmunoResearch) in 2% skim milk in TBS-T for 2 h at RT. The signal was visualized by chemiluminescence with Immobilon Forte Western HRP substrate (Millipore), and imaged with an LAS-3000UV Mini (Fujifilm). The chemiluminescence intensities of the bands were analyzed with ImageJ software. As a loading control, the membrane was stained with Ponceau S (Cell Signaling Technology).

For western blot analysis using Neuro2a cells, cells were scraped off the culture dish mechanically and homogenized in RIPA buffer. The same procedure as above was followed, with rabbit anti-itgb3 antibody (1:500, 13166; Cell Signaling Technology) and peroxidase-conjugated anti-rabbit IgG (1:10000, 711-035-152; Jackson ImmunoResearch) as primary and secondary antibody, respectively.

### Image analysis and quantification of cortico-mesencephalic axons

Images of labeled axons in the midbrain were obtained by confocal microscopy (ECLIPSE FN with EZ-C1; Nikon). For analysis, two consecutive 200-μm thickness coronal sections were used (Supplementary Figure 1). These sections are referred to as the anterior and posterior sections. For DiI-labeled samples, a total of 20–24 optical sections were acquired with a 10× objective lens (1024 × 1024 pixels; 1273 μm × 1273 μm) in 5-μm steps. For *in utero*-electroporated samples, 6–8 images were acquired using a 4× objective lens with a 2× digital zoom (1024 × 1024 pixels; 1591 μm × 1591 μm) in 20-μm steps. The projection images were thresholded using ImageJ (triangle method).

To evaluate the ectopic contralateral projection, the axon density index was defined by dividing the number of positive pixels on the contralateral (denervated) side by that of the ipsilateral (intact) side in each region of interest (ROI). For DiI-labeled samples, the size of the ROI was 400 × 1024 pixels (497 μm × 1273 μm), and the ROI was positioned at both sides adjacent to the midline. For EGFP-labeled samples, the size of the ROI on the ipsilateral side was 150 × 600 pixels (233 μm × 932 μm), which includes the EW. The size of the ROI on the contralateral side was 600 × 700 pixels (932 μm × 1088 μm), which contains a larger area of the denervated midbrain. A smaller ROI (450 × 450 pixels; 699 μm × 699 μm) was used to calculate the axon density index of the ventrolateral part of the denervated midbrain. To produce a pseudo-color heat-map image, the ROI of the contralateral side was divided into a grid pattern (12 × 14 grids, 50 × 50 pixels for each grid), and the axon density index was calculated for each grid. Samples in which axonal projection was rarely observed in the ipsilateral midbrain were excluded from the analysis.

### Statistical analysis

All statistical values are presented as the mean ± SEM. Statistical analyses were performed using the Mann–Whitney *U* test, the Student’s *t*-test, and one-way ANOVA with Tukey’s *post hoc* test. Differences between groups were considered to be significant at p < 0.05.

## Acknowledgements

We thank Dr. Ian Smith and Dr. Fujio Murakami for critical reading of the manuscript. This work was supported by MEXT KAKENHI on Dynamic regulation of brain function by Scrap & Build system (No. 16H06460) and JSPS KAKENHI Grant Nos. 19H03325 to N.Y., and 20J13844 to L.C. We would like to thank the Otsuka Toshimi Scholarship Foundation for scholarship support to L.C.

## References

Bareyre FM, Haudenschild B, Schwab ME (2002) Long-lasting sprouting and gene expression changes induced by the monoclonal antibody IN-1 in the adult spinal cord. J Neurosci 22:7097–7110.

Blackmore MG, Wang Z, Lerch JK, Motti D, Zhang YP, Shields CB, Lee JK, Goldberg JL, Lemmon VP, Bixby JL (2012) Krüppel-like Factor 7 engineered for transcriptional activation promotes axon regeneration in the adult corticospinal tract. Proc Natl Acad Sci 109:7517–7522.

Benowitz LI, Carmichael ST (2010) Promoting axonal rewiring to improve outcome after stroke. Neurobiol Dis 37:259–66.

Cafferty WBJ, Strittmatter SM (2006) The Nogo–Nogo receptor pathway limits a spectrum of adult CNS axonal growth. J Neurosci 26:12242–12250.

Carmel JB, Berrol LJ, Brus-Ramer M, Martin JH (2010) Chronic electrical stimulation of the intact corticospinal system after unilateral injury restores skilled locomotor control and promotes spinal axon outgrowth. J Neurosci 30:10918–10926.

Carmel JB, Martin JH (2014) Motor cortex electrical stimulation augments sprouting of the corticospinal tract and promotes recovery of motor function. Front Integr Neurosci 8:51.

Cohen-Cory S, Kidane AH, Shirkey NJ, Marshak S (2010) Brain-derived neurotrophic factor and the development of structural neuronal connectivity. Dev Neurobiol 70:271–288.

Cong L, Ran FA, Cox D, Lin S, Barretto R, Habib N, Hsu PD, Wu X, Jiang W, Marraffini LA, others (2013) Multiplex genome engineering using CRISPR/Cas systems. Science (80-) 339:819–823.

Dallner C, Woods AG, Deller T, Kirsch M, Hofmann H-D (2002) CNTF and CNTF receptor alpha are constitutively expressed by astrocytes in the mouse brain. Glia 37:374–378.

den Brand R, Heutschi J, Barraud Q, DiGiovanna J, Bartholdi K, Huerlimann M, Friedli L, Vollenweider I, Moraud EM, Duis S, others (2012) Restoring voluntary control of locomotion after paralyzing spinal cord injury. Science 336:1182–1185.

Doench JG, Fusi N, Sullender M, Hegde M, Vaimberg EW, Donovan KF, Smith I, Tothova Z, Wilen C, Orchard R, others (2016) Optimized sgRNA design to maximize activity and minimize off-target effects of CRISPR-Cas9. Nat Biotechnol 34:184–191.

Dougherty KD, Dreyfus CF, Black IB (2000) Brain-derived neurotrophic factor in astrocytes, oligodendrocytes, and microglia/macrophages after spinal cord injury. Neurobiol Dis 7:574–585.

Duan X, Qiao M, Bei F, Kim I-J, He Z, Sanes JR (2015) Subtype-specific regeneration of retinal ganglion cells following axotomy: effects of osteopontin and mTOR signaling. Neuron 85:1244–1256.

Ellison JA, Velier JJ, Spera P, Jonak ZL, Wang X, Barone FC, Feuerstein GZ (1998) Osteopontin and its integrin receptor aVb3 are upregulated during formation of the glial scar after focal stroke. Stroke-a J Cereb Circ 29:1698–1706.

Fink KL, López-Giráldez F, Kim I-J, Strittmatter SM, Cafferty WBJ (2017) Identification of intrinsic axon growth modulators for intact CNS neurons after injury. Cell Rep 18:2687–2701.

Fukuchi-Shimogori T, Grove EA (2001) Neocortex patterning by the secreted signaling molecule FGF8. Science 294:1071–1074.

Granseth B, Fukushima Y, Sugo N, Lagnado L, Yamamoto N (2013) Regulation of thalamocortical axon branching by BDNF and synaptic vesicle cycling. Front Neural Circuits 7:202.

Hatanaka Y, Murakami F (2002) In vitro analysis of the origin, migratory behavior, and maturation of cortical pyramidal cells. J Comp Neurol 454:1–14.

Hayano Y, Sasaki K, Ohmura N, Takemoto M, Maeda Y, Yamashita T, Hata Y, Kitada K, Yamamoto N (2014) Netrin-4 regulates thalamocortical axon branching in an activity-dependent fashion. Proc Natl Acad Sci 111:15226–15231.

Hu X, Leak RK, Shi Y, Suenaga J, Gao Y, Zheng P, Chen J (2015) Microglial and macrophage polarization—new prospects for brain repair. Nat Rev Neurol 11:56.

Humphries JD, Byron A, Humphries MJ (2006) Integrin ligands at a glance. J Cell Sci 119:3901–3903.

Kaiser J, Maibach M, Salpeter I, Hagenbuch N, de Souza VBC, Robinson MD, Schwab ME (2019) The spinal transcriptome after cortical stroke: in search of molecular factors regulating spontaneous recovery in the spinal cord. J Neurosci 39:4714–4726.

Ke M-T, Nakai Y, Fujimoto S, Takayama R, Yoshida S, Kitajima TS, Sato M, Imai T (2016) Super-resolution mapping of neuronal circuitry with an index-optimized clearing agent. Cell Rep 14:2718–2732.

Kitagawa H, Sugo N, Morimatsu M, Arai Y, Yanagida T, Yamamoto N (2017) Activity-dependent dynamics of the transcription factor of cAMP-response element binding protein in cortical neurons revealed by single-molecule imaging. J Neurosci 37:1–10.

Kojima M, Otabi H, Kumanogoh H, Toyoda A, Ikawa M, Okabe M and Mizui T (2020a) Reduction in BDNF from Inefficient Precursor Conversion Influences Nest Building and Promotes Depressive-Like Behavior in Mice. Int J Mol Sci 21: 3984.

Kojima M, Ishii C, Sano Y, Mizui T and Furuichi T (2020b) Journey of brain-derived neurotrophic factor: from intracellular trafficking to secretion. Cell Tissue Res: 1–10.

Komatsu Y, Watakabe A, Hashikawa T, Tochitani S, Yamamori T (2005) Retinol-binding protein gene is highly expressed in higher-order association areas of the primate neocortex. Cereb cortex 15:96–108.

Kuang RZ, Kalil K (1990) Specificity of corticospinal axon arbors sprouting into denervated contralateral spinal cord. J Comp Neurol 302:461–472.

Kullander K, Croll SD, Zimmer M, Pan L, McClain J, Hughes V, Zabski S, DeChiara TM, Klein R, Yancopoulos GD, others (2001) Ephrin-B3 is the midline barrier that prevents corticospinal tract axons from recrossing, allowing for unilateral motor control. Genes Dev 15:877–888.

Labandeira-Garcia JL, Costa-Besada MA, Labandeira CM, Villar-Cheda B, Rodríguez-Perez AI (2017) Insulin-like growth factor-1 and neuroinflammation. Front Aging Neurosci 9:365.

Labun K, Montague TG, Krause M, Torres Cleuren YN, Tjeldnes H, Valen E (2019) CHOPCHOP v3: expanding the CRISPR web toolbox beyond genome editing. Nucleic Acids Res 47:W171–W174.

Laemmli UK (1970) Cleavage of structural proteins during the assembly of the head of bacteriophage T4. Nature 227:680–685.

Lang C, Bradley PM, Jacobi A, Kerschensteiner M, Bareyre FM (2013) STAT3 promotes corticospinal remodelling and functional recovery after spinal cord injury. EMBO Rep 14:931–937.

Langmead B, Salzberg SL (2012) Fast gapped-read alignment with Bowtie 2. Nat Methods 9:357.

Lee J-K, Kim J-E, Sivula M, Strittmatter SM (2004) Nogo receptor antagonism promotes stroke recovery by enhancing axonal plasticity. J Neurosci 24:6209–6217.

Lein ES, Hawrylycz MJ, Ao N, Ayres M, Bensinger A, Bernard A, Boe AF, Boguski MS, Brockway KS, Byrnes EJ, others (2007) Genome-wide atlas of gene expression in the adult mouse brain. Nature 445:168–176.

Li B, Dewey CN (2011) RSEM: accurate transcript quantification from RNA-Seq data with or without a reference genome. BMC Bioinformatics 12:323.

Liang F, Hatanaka Y, Saito H, Yamamori T, Hashikawa T (2000) Differential expression of gamma-aminobutyric acid type B receptor-1a and-1b mRNA variants in GABA and non-GABAergic neurons of the rat brain. J Comp Neurol 416:475–495.

Liddelow SA, Barres BA (2017) Reactive astrocytes: production, function, and therapeutic potential. Immunity 46:957–967.

Luo W, Mizuno H, Iwata R, Nakazawa S, Yasuda K, Itohara S, Iwasato T (2016) Supernova: a versatile vector system for single-cell labeling and gene function studies in vivo. Sci Rep 6:35747.

Maddalo D, Manchado E, Concepcion CP, Bonetti C, Vidigal JA, Han Y-C, Ogrodowski P, Crippa A, Rekhtman N, de Stanchina E, others (2014) In vivo engineering of oncogenic chromosomal rearrangements with the CRISPR/Cas9 system. Nature 516:423–427.

McGraw J, Oschipok LW, Liu J, Hiebert GW, Mak CFW, Horie H, Kadoya T, Steeves JD, Ramer MS, Tetzlaff W (2004) Galectin-1 expression correlates with the regenerative potential of rubrospinal and spinal motoneurons. Neuroscience 128:713–719.

Mizuno H, Luo W, Tarusawa E, Saito YM, Sato T, Yoshimura Y, Itohara S, Iwasato T (2014) NMDAR-regulated dynamics of layer 4 neuronal dendrites during thalamocortical reorganization in neonates. Neuron 82:365–379.

Murakami F, Higashi S (1988) Presence of crossed corticorubral fibers and increase of crossed projections after unilateral lesions of the cerebral cortex of the kitten: a demonstration using anterograde transport of Phaseolus vulgaris leucoagglutinin. Brain Res 447:98–108.

Myers JP, Santiago-Medina M, Gomez TM (2011) Regulation of axonal outgrowth and pathfinding by integrin--ECM interactions. Dev Neurobiol 71:901–923.

Nah SH, Leong SK (1976) Bilateral corticofugal projection to the red nucleus after neonatal lesions in the albino rat. Brain Res 107:433–436.

Nieuwenhuis B, Haenzi B, Andrews MR, Verhaagen J, Fawcett JW (2018) Integrins promote axonal regeneration after injury of the nervous system. Biol Rev 93:1339–1362.

Nishitsuji K, Hosono T, Uchimura K, Michikawa M (2011) Lipoprotein lipase is a novel amyloid β (Aβ)-binding protein that promotes glycosaminoglycan-dependent cellular uptake of Aβ in astrocytes. J Biol Chem 286:6393–6401.

Omoto S, Ueno M, Mochio S, Takai T, Yamashita T (2010) Genetic deletion of paired immunoglobulin-like receptor B does not promote axonal plasticity or functional recovery after traumatic brain injury. J Neurosci 30:13045–13052.

Pang PT, Teng HK, Zaitsev E, Woo NT, Sakata K, Zhen S, Teng KK, Yung W-H, Hempstead BL, Lu B (2004) Cleavage of proBDNF by tPA/plasmin is essential for long-term hippocampal plasticity. Science 306:487–491.

Saito T, Nakatsuji N (2001) Efficient gene transfer into the embryonic mouse brain using in vivo electroporation. Dev Biol 240:237–246.

Sasaki K, Arimoto K, Kankawa K, Terada C, Yamamori T, Watakabe A, Yamamoto N (2020) Rho Guanine Nucleotide Exchange Factors Regulate Horizontal Axon Branching of Cortical Upper Layer Neurons. Cereb Cortex 30:2506–2518.

Serradj N, Paixão S, Sobocki T, Feinberg M, Klein R, Kullander K, & Martin JH (2014) EphA4-mediated ipsilateral corticospinal tract misprojections are necessary for bilateral voluntary movements but not bilateral stereotypic locomotion. J Neurosci 34: 5211–5221.

Silver J, Miller JH (2004) Regeneration beyond the glial scar. Nat Rev Neurosci 5:146–156.

Sinclair C, Mirakhur M, Kirk J, Farrell M, McQuaid S (2005) Up-regulation of osteopontin and αB-crystallin in the normal-appearing white matter of multiple sclerosis: an immunohistochemical study utilizing tissue microarrays. Neuropathol Appl Neurobiol 31:292–303.

Sirko S, Irmler M, Gascón S, Bek S, Schneider S, Dimou L, Obermann J, De Souza Paiva D, Poirier F, Beckers J, others (2015) Astrocyte reactivity after brain injury—: the role of galectins 1 and 3. Glia 63:2340–2361.

Starkey ML, Bartus K, Barritt AW, Bradbury EJ (2012) Chondroitinase ABC promotes compensatory sprouting of the intact corticospinal tract and recovery of forelimb function following unilateral pyramidotomy in adult mice. Eur J Neurosci 36:3665–3678.

Tabata H, Nakajima K (2001) Efficient *in utero* gene transfer system to the developing mouse brain using electroporation: visualization of neuronal migration in the developing cortex. Neuroscience 103:865–872.

Takahashi M, Vattanajun A, Umeda T, Isa K, Isa T (2009) Large-scale reorganization of corticofugal fibers after neonatal hemidecortication for functional restoration of forelimb movements. Eur J Neurosci 30:1878–1887.

Tom VJ, Doller CM, Malouf AT, Silver J (2004) Astrocyte-associated fibronectin is critical for axonal regeneration in adult white matter. J Neurosci 24:9282–9290.

Toshniwal PK, Firestone SL, Barlow GH, Tiku ML (1987) Characterization of astrocyte plasminogen activator. J Neurol Sci 80:277–287.

Tsirka SE, Gualandris A, Amaral DG, Strickland S (1995) Excitotoxin-induced neuronal degeneration and seizure are mediated by tissue plasminogen activator. Nature 377:340–344.

Tsujioka H, Yamashita T (2019) Comparison of gene expression profile of the spinal cord of sprouting-capable neonatal and sprouting-incapable adult mice. BMC Genomics 20:619.

Tsukahara N (1981) Synaptic plasticity in the mammalian central nervous system. Annu Rev Neurosci 4:351–379.

Ueno M, Hayano Y, Nakagawa H, Yamashita T (2012) Intraspinal rewiring of the corticospinal tract requires target-derived brain-derived neurotrophic factor and compensates lost function after brain injury. Brain 135:1253–1267.

Uesaka N, Hirai S, Maruyama T, Ruthazer ES, Yamamoto N (2005) Activity dependence of cortical axon branch formation: a morphological and electrophysiological study using organotypic slice cultures. J Neurosci 25:1–9.

Wahl A-S, Omlor W, Rubio JC, Chen JL, Zheng H, Schröter A, Gullo M, Weinmann O, Kobayashi K, Helmchen F, others (2014) Asynchronous therapy restores motor control by rewiring of the rat corticospinal tract after stroke. Science 344:1250–1255.

Waller R, Woodroofe MN, Wharton SB, Ince PG, Francese S, Heath PR, Cudzich-Madry A, Thomas RH, Rounding N, Sharrack B, others (2016) Gene expression profiling of the astrocyte transcriptome in multiple sclerosis normal appearing white matter reveals a neuroprotective role. J Neuroimmunol 299:139–146.

Wright MC, Mi R, Connor E, Reed N, Vyas A, Alspalter M, Coppola G, Geschwind DH, Brushart TM, Höke A (2014) Novel roles for osteopontin and clusterin in peripheral motor and sensory axon regeneration. J Neurosci 34:1689–1700.

Yamamoto N, Kurotani T, Toyama K (1989) Neural connections between the lateral geniculate nucleus and visual cortex in vitro. Science (80-) 245:192–194.

Yamamoto N, Yamada K, Kurotani T, Toyama K (1992) Laminar specificity of extrinsic cortical connections studied in coculture preparations. Neuron 9:217–228.

Yang J, Siao C-J, Nagappan G, Marinic T, Jing D, McGrath K, Chen Z-Y, Mark W, Tessarollo L, Lee FS, others (2009) Neuronal release of proBDNF. Nat Neurosci 12:113.

Yap E-L, Greenberg ME (2018) Activity-regulated transcription: bridging the gap between neural activity and behavior. Neuron 100:330–348.

Yokoyama N, Romero MI, Cowan CA, Galvan P, Helmbacher F, Charnay P, Parada LF, Henkemeyer M (2001) Forward signaling mediated by ephrin-B3 prevents contralateral corticospinal axons from recrossing the spinal cord midline. Neuron 29:85–97.

Zhang Y, Chen K, Sloan SA, Bennett ML, Scholze AR, O’Keeffe S, Phatnani HP, Guarnieri P, Caneda C, Ruderisch N, others (2014) An RNA-sequencing transcriptome and splicing database of glia, neurons, and vascular cells of the cerebral cortex. J Neurosci 34:11929–11947.

Zhong Y, Takemoto M, Fukuda T, Hattori Y, Murakami F, Nakajima D, Nakayama M, Yamamoto N (2004) Identification of the genes that are expressed in the upper layers of the neocortex. Cereb Cortex 14:1144–1152.

Zamanian JL, Xu L, Foo LC, Nouri N, Zhou L, Giffard RG, Barres BA (2012) Genomic analysis of reactive astrogliosis. Journal of neuroscience 32: 6391–6410.

