## Supplementary Information for "Involvement of denervated midbrain-derived factors in the formation of ectopic cortico-mesencephalic projection after hemispherectomy"

**Supplementary File 1.** List of differentially expressed genes (DEGs) in denervated midbrain at P8 and P10.

**Supplementary Tables**: 2

**Supplementary Figures**: 7

**Supplementary Table 1.** Primers used for ISH probe synthesis.

| Gene symbol | Forward | Reverse |
|---|---|---|
| Spp1 | AATCTCCTTGCGCCACAG | TGGCCGTTTGCATTTCTT |
| Vim | CCTCATTCCCTTGTTGCAGTT | AGTGAGGTCAGGCTTGGAAAC |
| Abca1 | GAAGGACCACTAAGTAACCCCC | GGCTGTACATGGTGATAATGGA |
| Tyrobp | TGGACTGTGGTGTCCAGTG | TTCCAACGGGAGTCCTGA |
| Fcgr3 | TGCACACTCTGGAAGCCA | AGAACCAAACATCCCGGC |
| Plat | CTCACGTCAGACTGTACCCG | GACCAGGAGGGCAGACTTTG |
| Fn1 | CCCTTACAGTTCCAAGTTCCTG | AAAGGCTTAAGGGTGAAAGGAC |

**Supplementary Table 2.** sgRNA sequences for each candidate receptor.

| **Gene symbol** | **sgRNA target sequence** | |
|---|---|---|
| | Forward | Reverse |
| **Cd44** | GCAATATGTGTCATAGTGGG | CAGTCCGGGAGATACTGTAG |
| **Plaur** | AGGACCATGAGTTACCGCAT | AGGTGCAGGACGCACACTCG |
| **Notch1** | ATATACATACCCGTCCACTC | ACCTGCAAGGACATGACCAG |
| **Cntfr** | ATACTGCGAAGCTTAGAACT | GGCCGTAGTGTTGTGACCCA |
| **Itgb1** | GAGGAATGTAACACGACTGC | TCCCAACATTCCTACCAATG |
| **Itgb3** | GCAGGTGGAGGATTACCCCG | AATATGGGTCTTGGCATCCG |
| **Tgfbr1** | AGAGCGTTCATGGTTCCGAG | ATGAAAGGGCGATCTAGTGA |
| **Vldlr** | TAGAATACAAACCCCGACAA | AGTGCATTCACAAAAAATGG |
| **Lrp8 (Apoer2)** | CGATGGCTCTGACGAATCGA | TCGCAGGGGGTCAACGGCAA |
| **Lrp1** | GCTGCGGAACAGTACCACGT | ACTGCGTCTTACCCAGACTG |
| **Igf1r** | AGAACCGAATCATCATAACG | CTGCTTATTAACATCCGGAG |
| **Ntrk2 (TrkB)** | ATGACGTTGAAGCTTACGTG | AACCTGCAGATACCCAATTG |

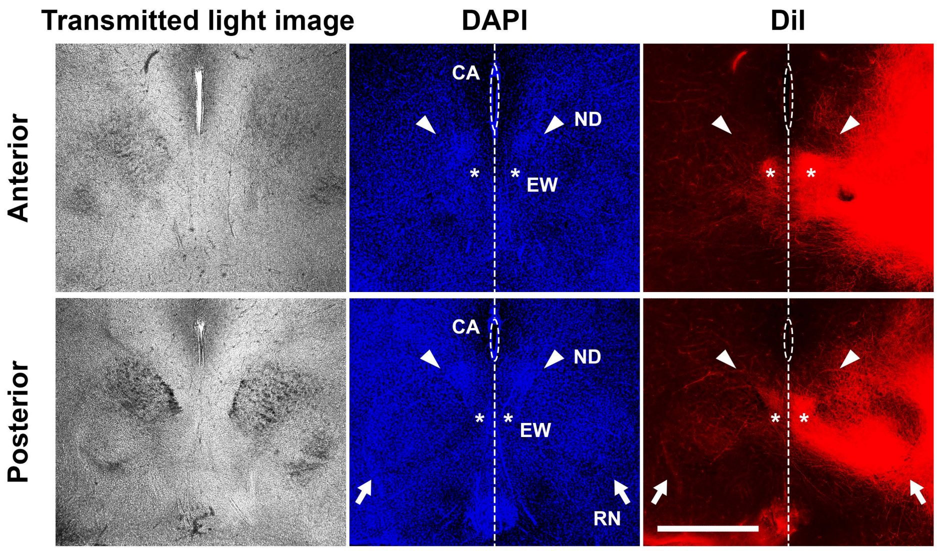


**Supplementary Figure 1.** Serial sections of the midbrain show DiI-labeled contralateral axons in the midbrain with hemispherectomy. Asterisks and arrows indicate the EW and RN, respectively. CA; cerebral aqueduct, ND; nucleus of darkschewitsch (arrowheads). Note that axons accumulate near the midline (dashed lines) in the region ventral to the ND, around the EW. Scale bar = 1 mm.

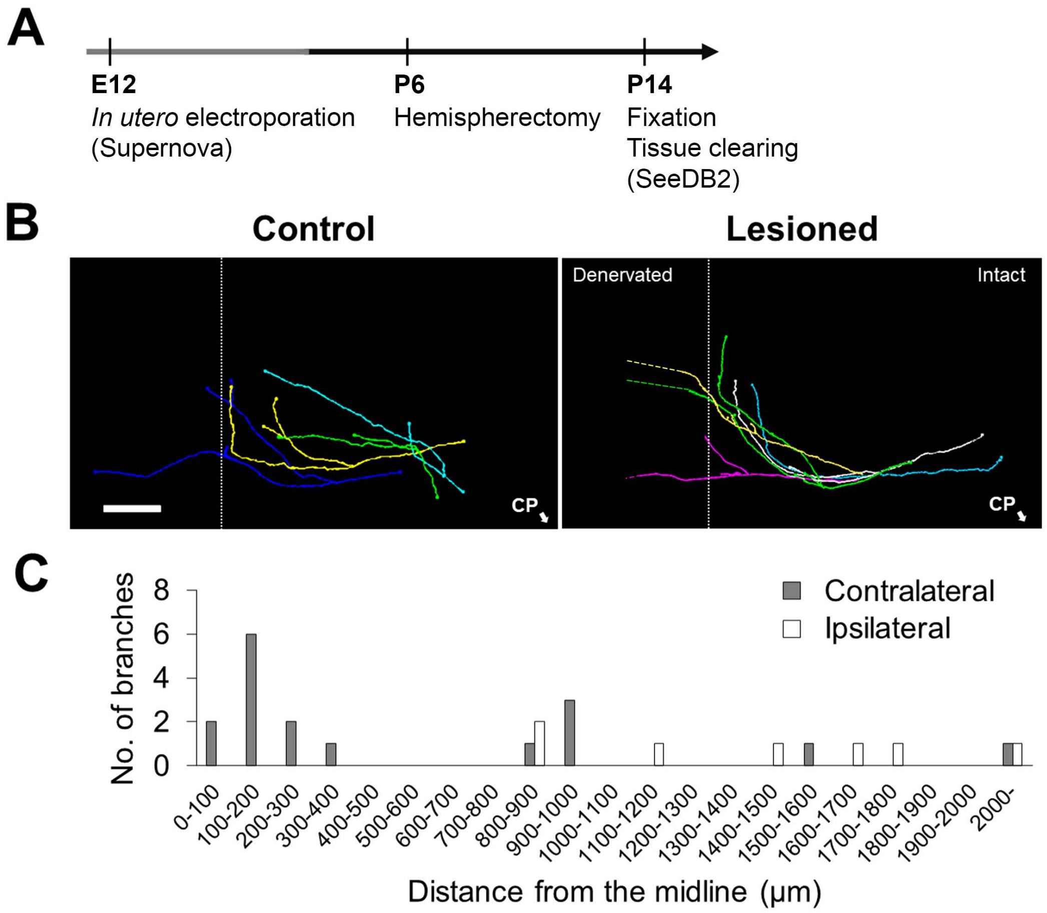


**Supplementary Figure 2.** Axonal sprouting occurs in the region adjacent to the midline to project contralaterally. **A,** Schematic overview of the experiment. Layer 5 neurons were sparsely labeled by the Supernova method. Tissue clearing was performed in thick (1-mm thickness) midbrain sections to trace labeled axons. **B,** Representative axon trace images of control and hemispherectomized samples. Branches of contralaterally projecting axons formed in the region adjacent to the midline (arrowheads). White lines indicate the midline, and the location of the CP is indicated by the arrows. Scale bar = 500 μm. **C,** Histogram showing the distribution of the distance between midline and branch points. Note that contralaterally projecting axons form branches within 300 μm of the midline, while both contralaterally and ipsilaterally projecting axons have branches in the region more than 800 μm away from the midline (n = 7 axons for each group).

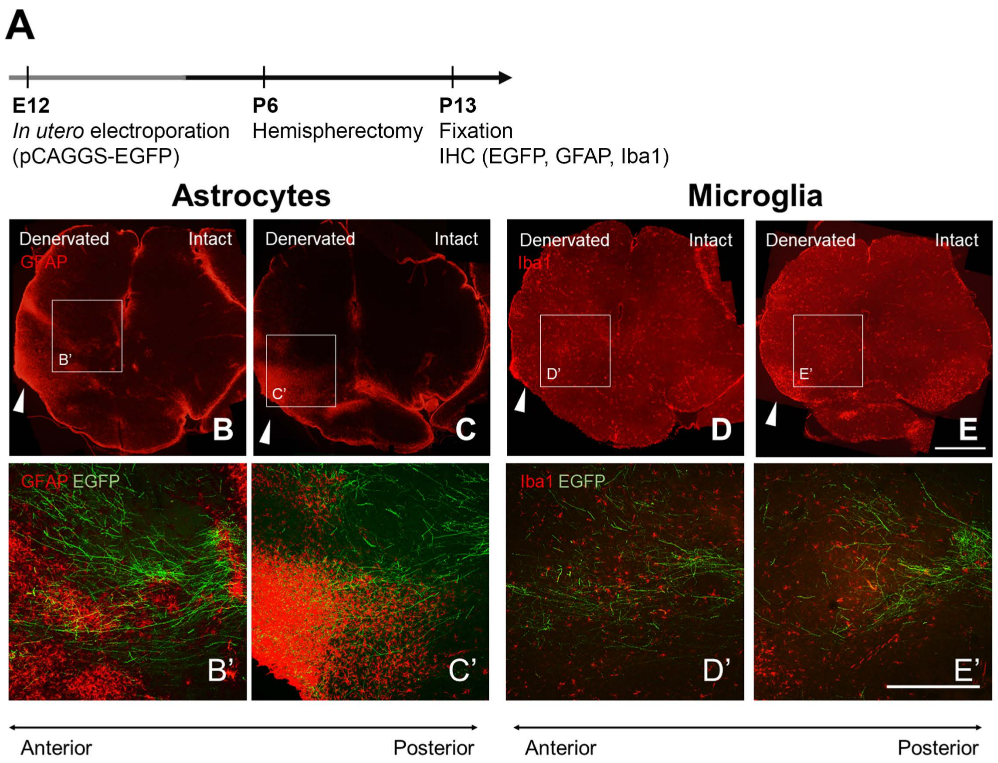


**Supplementary Figure 3.** Distribution of glial cells in the midbrain after hemispherectomy. **A,** Schematic overview of the experiment. Immunohistochemistry with glial cell markers (GFAP and Iba1) was performed after hemispherectomy. *In utero* electroporation was conducted as described previously to label corticofugal axons. **B–E,** Distribution of GFAP-positive reactive astrocytes (B, C) and Iba1-positive microglia (D, E) along the anterior–posterior axis. Lower panels are high-magnification views of the boxed areas above. Scale bar = 1 mm (upper panels), 500 µm (lower panels). Strong GFAP signals were observed near the CP on the denervated side (arrowheads), with broad distribution across the ventral part of the midbrain (B, C). Strong Iba1-positive signals near the denervated CP had ameboid morphology. Microglia distant from the CP had ramified morphology (D, E). Many contralaterally projecting axons run near GFAP-positive cells (B', C') and Iba1-positive cells (D', E').

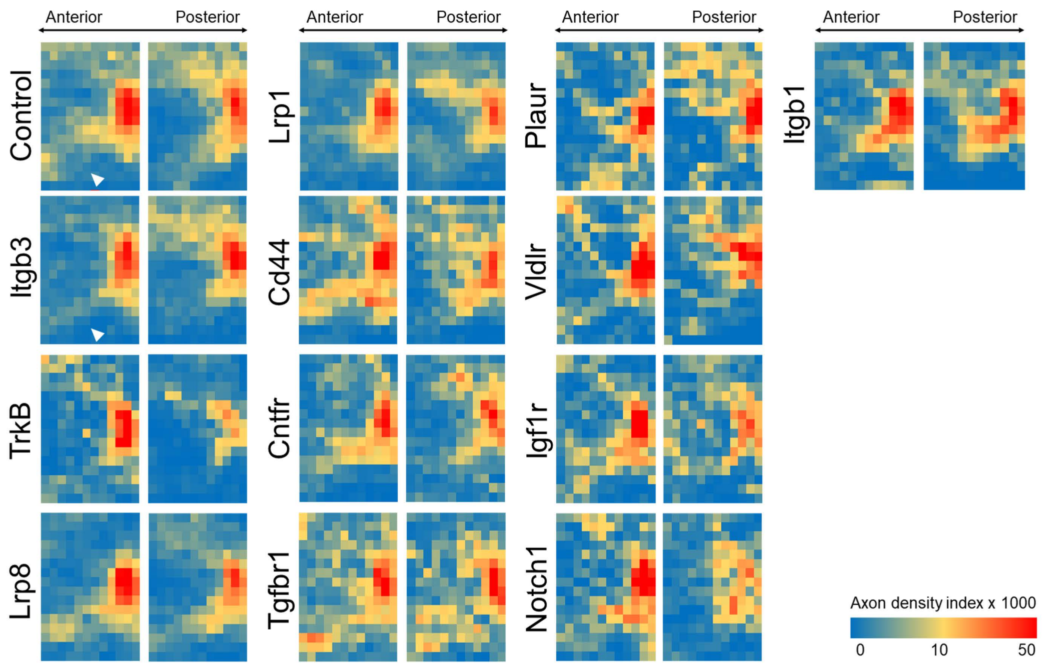


**Supplementary Figure 4.** Axon density indices in the denervated midbrain of the control and 12 candidate receptor-KOs after hemispherectomy are shown as pseudo-color heat-map images. Note that the axon density indices in the ventrolateral portion of the anterior side are greatly reduced in the Itgb3 KO sample (arrowheads), and overall axon density indices are reduced in the TrkB KO sample.

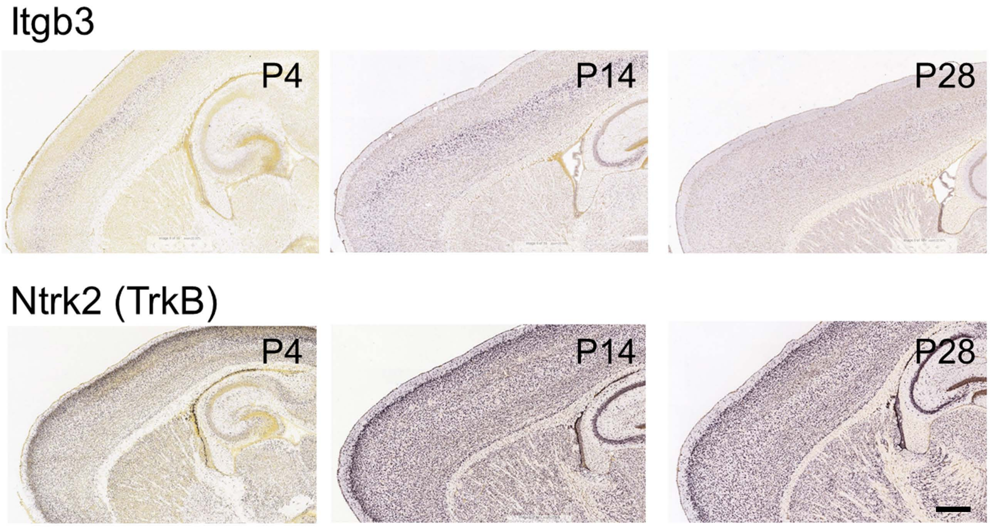


**Supplementary Figure 5.** Expression of Itgb3 and TrkB in cortex. Itgb3 and TrkB were expressed in layer 5 cortical neurons in the Allen Developing Mouse Brain Atlas (*In situ* hybridization), especially in young mice up to 2 weeks old. Sagittal sections are shown. Image credit: Allen Institute (Itgb3: http://developingmouse.brain-map.org/gene/show/16189; Ntrk2: http://developingmouse.brain-ap.org/gene/show/17979). Scale bar = 500 µm.

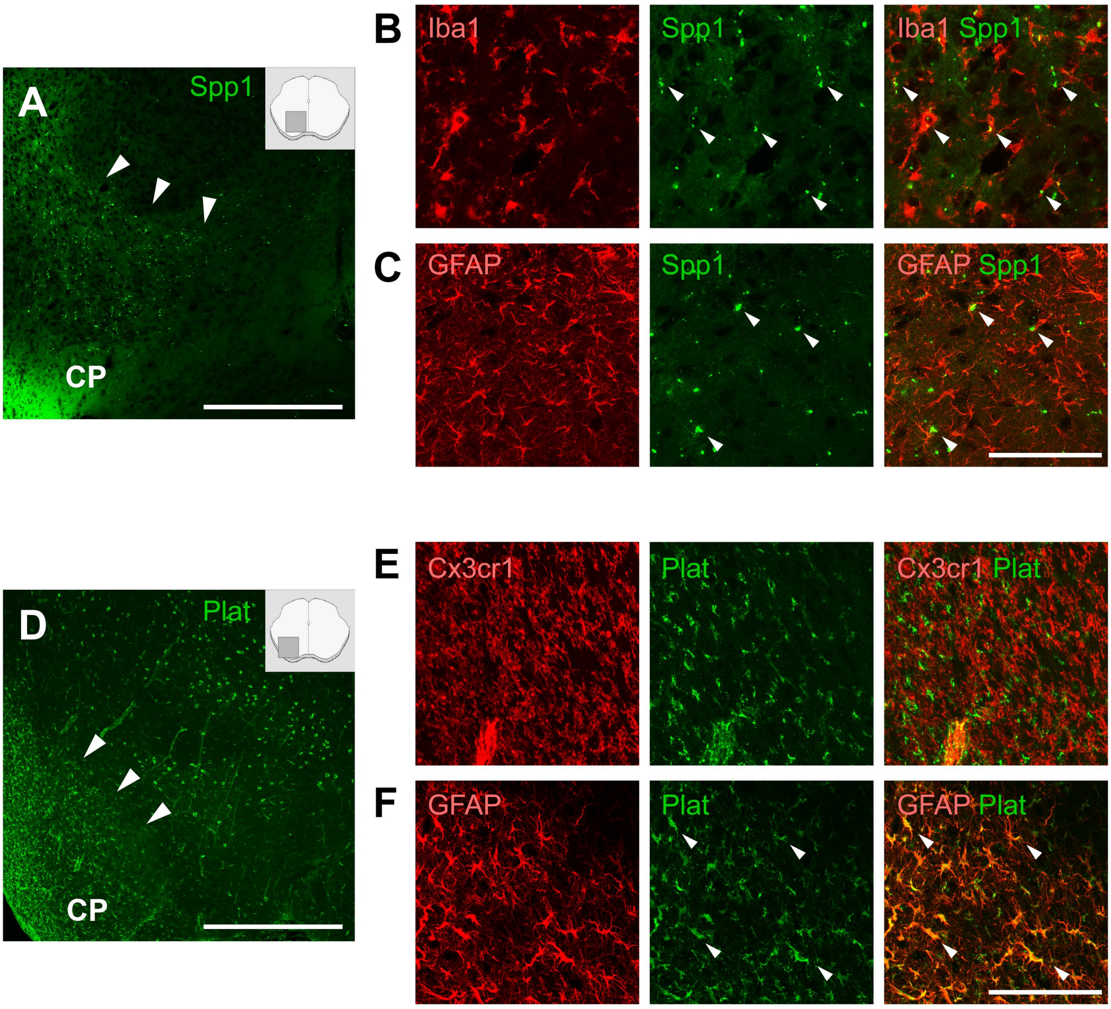


**Supplementary Figure 6.** Distribution of Spp1 (Osteopontin) and Plat (Tissue-type plasminogen activator) protein. **A,** Punctate signals of Spp1 were observed broadly on the ventral side of the denervated midbrain (arrowheads), with strong signals on the ventrolateral side near the CP. Illustration (insert) indicates the location of the imaged area. Scale bar = 500 µm. **B,** Immunostaining of Iba1 and Spp1 shows that Spp1 was detectable within the Iba1-positive microglia (arrowheads). **C,** Immunostaining of GFAP and Spp1 showed that Spp1-positive puncta reside in and around GFAP-positive astrocytes (arrowheads). Scale bar = 100 µm. **D,** Immunopositive signals for Plat were observed on the area near the CP (arrowheads). Scale bar = 500 µm. **E,** Immunostaining of Cx3cr1 and Plat shows that Plat signals do not co-localize with Cx3cr1-positive microglia. **F,** Immunostaining of GFAP and Plat showed that Plat-positive signals co-localize with GFAP-positive astrocytes (arrowheads). Scale bar = 100 µm.

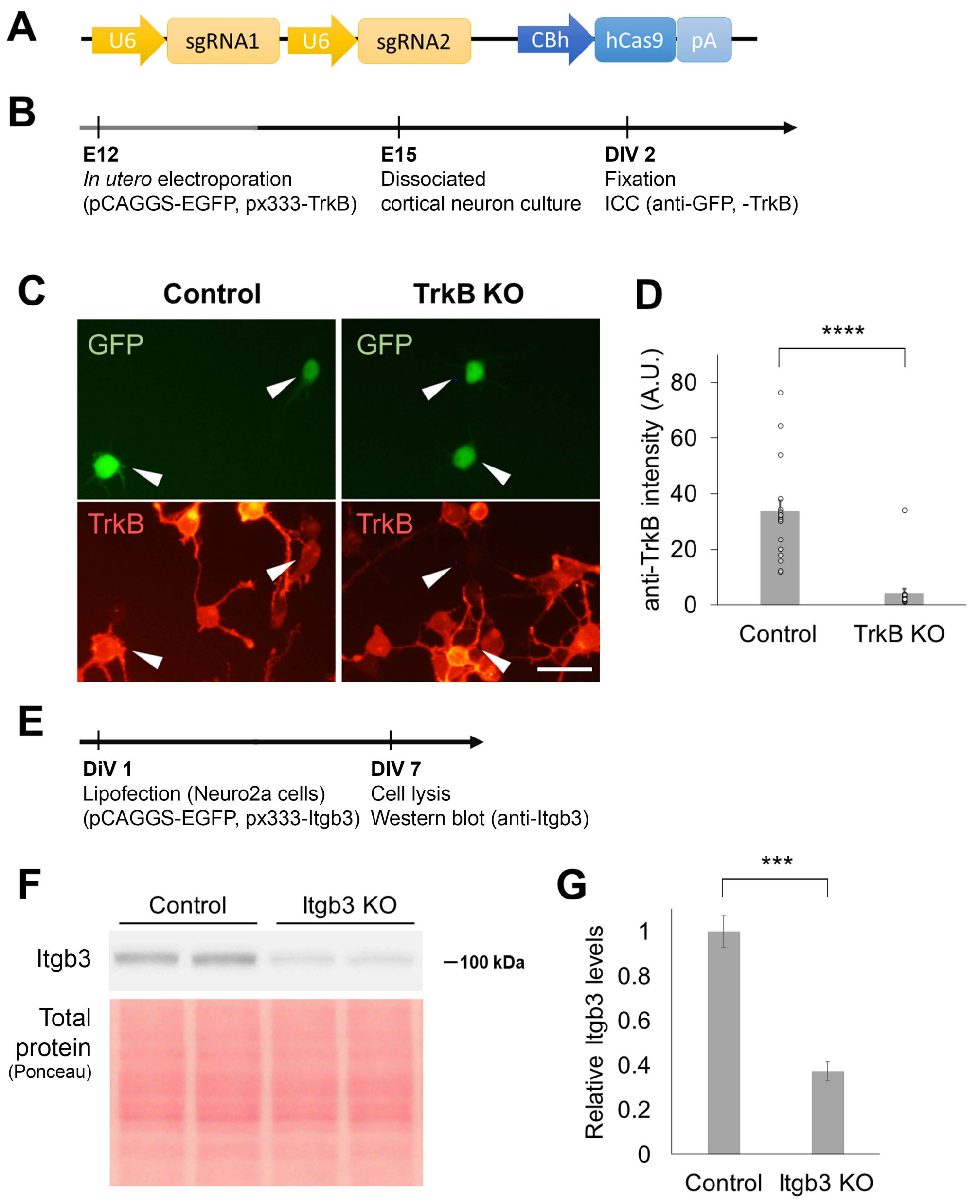


**Supplementary Figure 7.** Validation of CRISPR/Cas9-mediated KO efficiency. **A,** A plasmid expressing dual single-guide RNAs and hCas9 (shown) was transfected into the cortex together with an EGFP-expressing plasmid to knock out TrkB. **B,** Schematic overview of the experiment to validate TrkB KO. Dissociated neuron cultures were prepared from cortex transfected with the KO plasmid and were subjected to immunocytochemistry (ICC) with anti-TrkB. **C,** Most GFP-positive neurons were TrkB-negative (arrowheads). Scale bar = 30 μm. **D,** Immunostaining intensity was considerably reduced by TrkB KO (Control, 33.8 ± 3.9, n = 18; TrkB KO, 4.2 ± 1.6, n = 19). **** $p < 0.0001$, Student's *t*-test. **E,** Schematic overview of the experiment to validate Itgb3 KO. The plasmid for Itgb3 KO was transfected with the EGFP-expressing plasmid into Neuro2a cells by lipofection, and the cultured cells were subjected to western blot analysis. **F-G,** Itgb3 expression level was markedly decreased by Itgb3 KO (ratio, 0.37 ± 0.042 against control, n = 4). As the transfection efficiency of lipofection was 66.5 ± 4.0% (n = 4), the KO efficiency is estimated as 94.3%. *** $p < 0.001$, Student's *t*-test.
